# Activation of cGAS-STING signaling pathway during HCV infection

**DOI:** 10.1101/2025.01.11.632524

**Authors:** Saleem Anwar, Khursheed Ul Islam, Iqbal Azmi, Jawed Iqbal

**Affiliations:** Multidisciplinary Centre for Advanced Research and Studies, Jamia Millia Islamia, Jamia Nagar, New Delhi 110025, India

**Keywords:** HCV, cGAS, STING, IFN-β, cGAMP, RIG-I

## Abstract

Hepatitis C virus (HCV) is a major global health issue, infecting over 170 million people worldwide and leading to severe liver diseases, including cirrhosis and hepatocellular carcinoma. The ability of HCV to persist and cause chronic infection is partly due to its evasion of the host’s innate immune responses, particularly those mediated by the RIG-I-MAVS pathway, which is critical for antiviral defense. Studies have shown the crucial role genome sensing of DNA viruses by cyclic GMP-AMP synthase (cGAS) followed cGAMP production and activation of downstream effector STING (stimulator of interferon genes) to induce IFN-β, however it is not understood in RNA viruses specially in HCV infection. In this study, we explored first time the mechanism of the cGAS-STING pathway in the context of HCV infection, specifically using the JFH-1 strain of HCV (genotype 2a). We observed that cGAS expression is significantly upregulated during the early HCV infection, leading to the production of the second messenger cyclic GMP-AMP (cGAMP), which in turn activates STING. This activation results in the significant induction of type I interferon responses, particularly interferon-β (IFN-β), which is essential for mounting an effective antiviral response. Moreover, our results demonstrated the translocation of cGAS and STING with cellular organelles such as the endoplasmic reticulum (ER) and mitochondria. This suggests that the cGAS-STING pathway is intricately linked with other cellular signaling networks in detecting and responding to HCV infection. Furthermore, knockdown experiments targeting cGAS, STING and RIG-I revealed that these proteins play a crucial role in suppressing HCV replication, underscoring their potential as therapeutic targets. These findings provide valuable insights into the molecular mechanisms of the cGAS-STING pathway in mediating the innate immune response against HCV. Understanding this pathway’s role in the immune defense against HCV opens up new possibilities for therapeutic strategies aimed at enhancing the host’s antiviral immunity and potentially developing new treatments for chronic HCV infection.

## Introduction

Hepatitis C virus (HCV), a member of the Flaviviridae family, infects approximately 3% of the global population, which translates to around 170 million people (1). Hepatitis C remains a major public health issue, characterized by severe liver diseases such as cirrhosis and hepatocellular carcinoma. Due to its high mutation rate, HCV continuously evolves and is known to have seven major genotypes and over 67 subtypes (2,3). In India, HCV contributes significantly to the annual death toll. HCV contains a single-stranded positive-sense RNA genome, approximately 9.6 kb in length, is flanked by conserved, highly conserved non-translated regions (NTRs) essential for RNA translation and replication (4). The 5’ NTR contains an internal ribosome entry site (IRES) that regulates translation initiation, influenced by various cellular factors interacting with either the IRES sequence or the core encoding sequences. The polyprotein precursor encoded by the HCV genome, approximately 3,000 amino acids long, is processed by viral and cellular proteases into several proteins. The N-terminal region produces the structural core (C) and envelope proteins 1 and 2 (E1 and E2), crucial for forming infectious virus particles (5). C-terminal to E2 is the hydrophobic p7 protein, necessary for the efficient assembly and release of virions. The remaining polyprotein is cleaved into nonstructural (NS) proteins such as, NS2, NS3, NS4A, NS4B, NS5A, and NS5B (6). Due to the lack of a 3’-5’ exonuclease proofreading activity in the NS5B RNA-dependent RNA polymerase (RdRp), HCV RNA replication is error-prone, resulting in high genetic variability (7). Extensive research over the past two decades has significantly advanced our understanding of HCV replication, leading to the development of direct-acting antivirals (DAAs) (8). These DAAs are substantially effective against HCV genotype 1. However, the genetic distinction among all HCV genotypes can impact the efficacy of these treatments, necessitating continued research to develop therapies effective against all genotypes (9).

Hepatitis C virus (HCV) encodes several proteins that specifically target and suppress the host antiviral pathways, particularly those involving type-I interferons (IFNs). The NS3/4A protease cleaves and inactivates adaptor proteins like Mitochondrial antiviral-signaling protein (MAVS) and Toll/interleukin-1 receptor (TRIF), disrupting RIG-I-like receptor signaling crucial for IFN induction (10). The non-structural protein (NS5A) interacts with host factors to inhibit protein kinase R (PKR), reducing the ability of host to inhibit viral replication (11). The core protein interferes with the IFN signaling pathway by modulating STAT1 activity (12), while the E2 glycoprotein binds to CD81 on host cells, inhibiting natural killer (NK) cell activity and weakening the innate immune response (13). NS4B possesses NTPase and RNA-binding activities, it regulates RNA-dependent RNA polymerase (RdRP) activity of NS5B, it also impacts endoplasmic reticulum function, and modulates both viral and host translation processes. (13). The RNA-dependent RNA polymerase (NS5B) is essential for viral RNA replication and interacts with host factors influencing the antiviral response (14). These HCV proteins collectively contribute to the suppression of the host antiviral defense mechanisms, facilitating viral replication and persistence in the liver. Understanding these interactions is crucial for developing targeted therapies to combat HCV infection effectively. To counteract these viral strategies, the host employs its own defense mechanisms, with innate immunity playing a crucial role in defending against microbial infections and initiating adaptive immune responses. Pattern recognition receptors (PRRs) such as Toll-like receptors (TLRs), RIG-I-like receptors (RLRs), Nod-like receptors (NLRs), and DNA sensors such as cGAS like receptors (cGLRs) and IFI16 detect pathogen-associated molecular patterns (PAMPs) of invasive microbes (15). Upon detection, PRRs trigger intracellular signaling cascades, leading to the induction of type I interferons and pro-inflammatory cytokines (16). The retinoic acid-inducible gene 1 (RIG-I) protein, encoded by the DDX58 gene, functions as a PRR in RNA virus infections by specifically binding to short dsRNA with a 5’-triphosphate. Normally, RIG-I is in a closed, auto-inhibited state. Binding of 5’-ppp-RNA to the C-terminal domain/helicase region activates RIG-I, promoting oligomerization and subsequent downstream signaling, including interaction with mitochondrial antiviral signaling protein (MAVS), the adaptor protein anchored to the mitochondrial outer membrane (17).

Building on previous studies demonstrating the activation of cGAS and STING during infections by various RNA viruses, and the observed crosstalk between the RIG-I-MAVS and cGAS-STING signaling pathways during HCV infection, our research aims to elucidate whether the activation of cGAS or STING is a direct response to the HCV genome or an indirect result of other mechanisms. Our findings reveal that both cGAS and STING are activated early in HCV infection, along with downstream signaling molecules (cGAMP, IFN-β etc). We observed an interaction between cGAS and dsRNA shortly after HCV infection, indicating a potential direct recognition mechanism. Additionally, we detected the expression of cGAMP following HCV infection, supporting the activation of the cGAS-STING pathway. Knockdown studies in our research demonstrated that the absence of cGAS or STING significantly impacts HCV replication and the IFN-β response. This suggests that both cGAS and STING play crucial roles in the antiviral defense against HCV. Our study adds first time to the understanding of the molecular mechanisms of cGAS-STING pathway activation during HCV infection.

## Results

### Verification of HCV infection in human hepatoma cells

HCV particles were generated and quantified using a previously established protocol (18). Briefly, the J6/JFH-1 (genotype 2a) RNA was *in vitro* transcribed and electroporated into Huh7.5 cells using Gene Pulser Xcell Total system (BioRad). The generation of HCV viral particles was confirmed by TaqMan probe based RTq-PCR using HCV gene-specific primers targeting the 5’ UTR region of the HCV genome (data not shown). The released virus (J6/JFH1) was quantified by FFU (data not shown) method as shown earlier (18) and (multiplicity of infection) MOI 1 was used for further infection of naïve Huh 7 cells. In contrast, supernatant of JFH-1/GND was incubated with Huh 7 cells which serves as mock or uninfected cells. To determine the HCV infection, total RNA was isolated from Huh7 cells infected with J6/JFH-1 and JFH-1/GND (replication defective) using the Trizol method as per the manufacturer’s protocol. After RNA isolation, cDNA synthesis was performed, followed by quantitative PCR (qPCR) to measure the levels of HCV gene expression. The presence of HCV RNA in J6/JFH-1 infected Huh7 cells confirmed successful replication and transcription of HCV. However, no virus was detected in JFH-1/GND incubated cells (mock) (Supplementary Fig. 1a).

To further confirm HCV infection in Huh7 cells, a western blot analysis was conducted. The above infected Huh7 cells were grown till day 3 post infection and cellular lysates were prepared and analyzed by western blotting. The results showed a marked expression of the HCV core protein in J6/JFH-1-infected cells compared to mock-infected controls (JFH1/GND). Actin was used as a protein loading control (Supplementary Fig. 1b).

### Induction of cGAS and STING expression during HCV infection

It is well established that during replication, HCV produces dsRNA intermediates in the cytoplasm of infected cells. These intermediates bind to RIG-I, which then triggers the activation of the mitochondrial antiviral signaling protein (MAVS) on the mitochondrial surface, leading to an antiviral response in the infected cells (17). Previous studies have shown that the HCV NS3/4A protease cleaves MAVS at Cys-508, leading to its dislocation from the mitochondria and thereby inhibiting IFN-β induction at later stage of infection (data not shown), disrupting the RIG-I-MAVS-mediated antiviral signaling (19). Given this, we sought to identify any parallel active pathways during this time point of infection. Considering the roles of cGAS and STING in many RNA viruses and the targeting of STING by HCV NS4B to prevent the formation of the RIG-I-MAVS-STING complex for interferon production, we aimed to elucidate the role of the cGAS-STING pathway during HCV infection. cGAS is well known to sense viral DNA and produces cGAMP by utilizing GTP and ATP (20). This cGAMP binds to STING and stimulate STING-IRF3 antiviral pathway to curb viral proliferation (17). However, cGAS sensing to viral RNA is not known yet but cGAS binding has been shown with a 50 mer RNA, however this binding is not enough to produce cGMAP (21). Additionally, a recent report revealed that a cGAS-like receptor (cGLR) in Drosophila binds to dsRNA. This binding triggers the production of cGAMP, which subsequently binds to and activates STING. The activated STING then stimulates downstream molecules to produce interferons, thereby initiating an antiviral response (22). Based on these observations, we sought to determine the status of cGAS during HCV infection. Total extracted RNA at early time points of HCV infection was used to analyze cGAS mRNA expression using its specific primers by RT-qPCR. The results show the significant induction of cGAS at 12h which was reduced at day1 infection **(Fig. 1a)**. However, enough cGAS induction was observed at 2h of HCV infection as well **(Fig. 1a)**. Together, this observation indicates the induction of cGAS at early HCV infection. Studies have shown that cGAS binds to viral nucleic acids leading to the production of cGAMP. This cGAMP binds to and activate STING, which further recruits and activates several other proteins, such as TBK1 and IRF3, to initiate an antiviral pathway (23). Since we observed cGAS induction during HCV infection, we wanted to investigate the status of STING expression. Further, total extracted RNA was utilized for RT-qPCR using STING specific primers. We observed some induction of STING at 2h, but significant induction at 12h and at day1 infection **(Fig. 1b)**.

**Figure 1.**
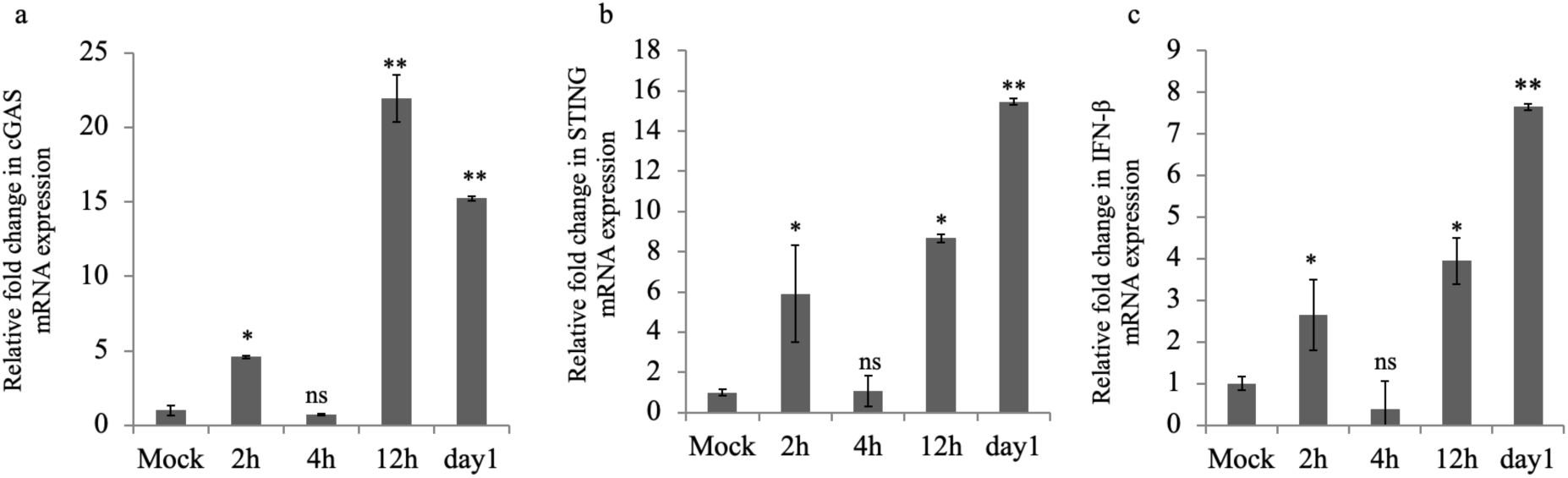
HCV induces cGAS and STING expression at early infection. (a-c) depicts the relative fold change in cGAS, STING and IFN-β mRNA expression at 2h, 12h, day1 of HCV infection. The values represent the means ± SDs from two independent experiments performed in duplicate. * p-value ≤ 0.05, ** p-value ≤0.01, ns (non significant), mock vs infection.

### Induction of interferon expression during HCV infection

Next, we examined the expression of IFN-β and observed a significant induction at 2 hours, 12 hours, and 1-day post-infection **(Fig. 1c)**. These findings suggest that activated STING may have recruited other signaling proteins, ultimately leading to IFN-β induction. Additionally, we assessed the early expression of IFN-α and IFN-γ during HCV infection, as both cytokines play key roles in the antiviral response against viral infections (24). Our observations revealed no difference in IFN-α expression between uninfected and HCV-infected cells at the early stage of infection (Supplementary Fig. 2). This suggests that IFN-α, despite its importance in immune responses to viruses, it is not induced for immediate immune response during the initial phase of HCV infection, or it is possible that Huh 7 cells do not express this as it is an epithelial origin cell line. However, we could not get any amplification of IFN-γ. Together, we observed a significant induction of IFN-β but no change in IFN-α expression and no amplification of IFN-γ gene was observed.

### cGAS translocation to endoplasmic reticulum and mitochondria during HCV infection

Previous studies have shown that HCV genomic RNA replication occurs within specialized replication organelles (RO), which are closely associated with modifications in the endoplasmic reticulum (ER) membrane. These organelles house viral replication complexes consisting of the non-structural proteins NS3 to NS5B (25). It has also been shown that STING is localized to the endoplasmic reticulum (ER), particularly within ER-mitochondria associated membranes (MAM) (26). Certain viral infections can induce ER stress, leading to the unfolded protein response (UPR), which in turn may alter the localization of various proteins (27) (28). Mitochondria function as key signaling hubs that regulate various cellular processes, particularly innate immune responses. They provide a scaffold for signaling molecules, enabling the assembly of signal complexes that drive immune activation. For example, the aggregation of the antiviral signaling protein MAVS on the mitochondrial outer membrane forms a critical platform for initiating the antiviral response (29).

Since, many proteins specially STING has been shown to be translocated to different organelles for its function (30), we sought to determine whether cGAS co-localizes with the ER and mitochondria at 12 h HCV infection when cGAS induction was observed at peak. To investigate this, we performed immunofluorescence studies using mock and HCV-infected cells with specific antibodies targeting cGAS, PD1 (ER marker), and TOM20 (mitochondrial marker). This allowed us to visualize and analyze the spatial distribution of cGAS with these organelles. Our findings demonstrated a significant co-localization of cGAS with endoplasmic reticulum but no colocalization with mitochondria in HCV-infected cells (**Fig. 2**).

**Figure 2.**
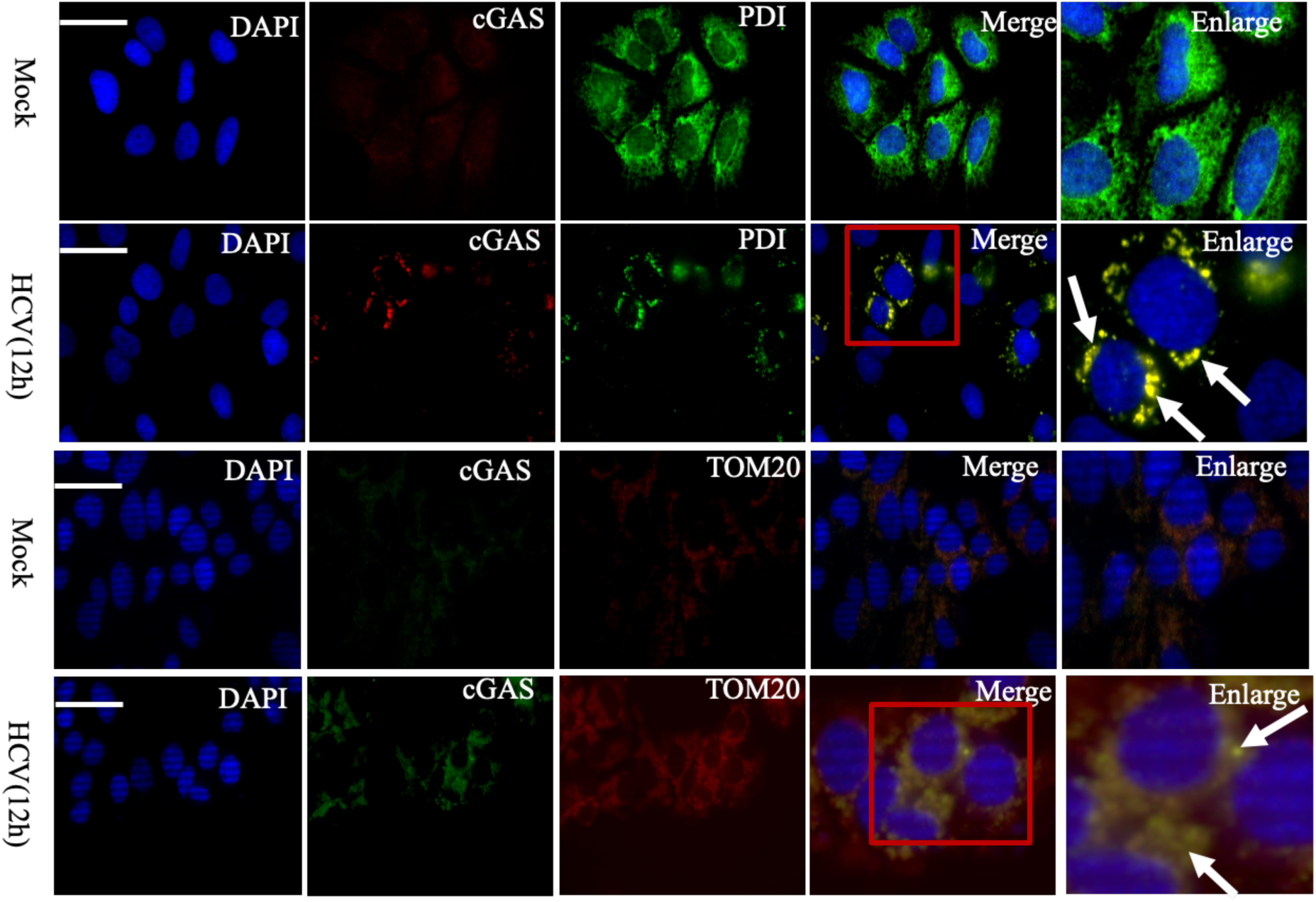
cGAS associates with endoplasmic reticulum and mitochondria at early infection. Huh 7 (mock) and HCV infected cells were fixed, permeabilized and stained with anti-cGAS (mouse), anti-PDI (rabbit) for cGAS-ER combination, and anti-TOM-20 (mouse) and anti-cGAS (rabbit) primary antibodies for cGAS-Mito combination. After washing with PBS, secondary antibodies Alexa Fluor-594 conjugated goat anti-mouse (red), and AlexaFluor-488 conjugated goat anti-rabbit (green) were used to study colocalization of cGAS with endoplasmic reticulum and mitochondria. DAPI was used to stain nuclei. Yellow dotes indicated by white arrows in the merge picture depicts the co-localization of two proteins. Images were captured at 63x magnification with a scale bar of 20μm under fluorescence microscope (Zeiss Axio observer 7).

### STING distribution to ER and mitochondria during HCV infection

STING is an endoplasmic reticulum (ER) resident protein that, upon binding to cGAMP, translocates from the ER to the Golgi. In the Golgi, STING initiates a type I interferon response by activating and phosphorylating IRF3 <u>(1).</u> STING plays an unexpected role in responding to RNA viruses, with mitochondria mediating the interaction (26). Since STING is an ER protein, and mitochondria is believed to be a platform for antiviral signaling pathway (31), We first aimed to assess STING colocalization with the ER, followed by determining whether activated STING colocalizes with mitochondria, where MAVS proteins are present, to mediate a type I interferon response. Huh7 and HCV-infected cells were fixed and permeabilized at 12 hours post-infection, then immunostained with antibodies against STING, PDI (ER marker), and TOM20 (mitochondria marker). The results show STING colocalization with both the ER and mitochondria, suggesting that for antiviral response STING may require its presence at the mitochondria, similar to its known association with the Golgi and perinuclear bodies (30). This translocation is crucial for activating downstream signaling pathways that trigger interferon and cytokine production (23). STING localization with ER is reduced after infection that could be due to its translocation to other organelles (**Fig. 3**), which require further study.

**Figure 3.**
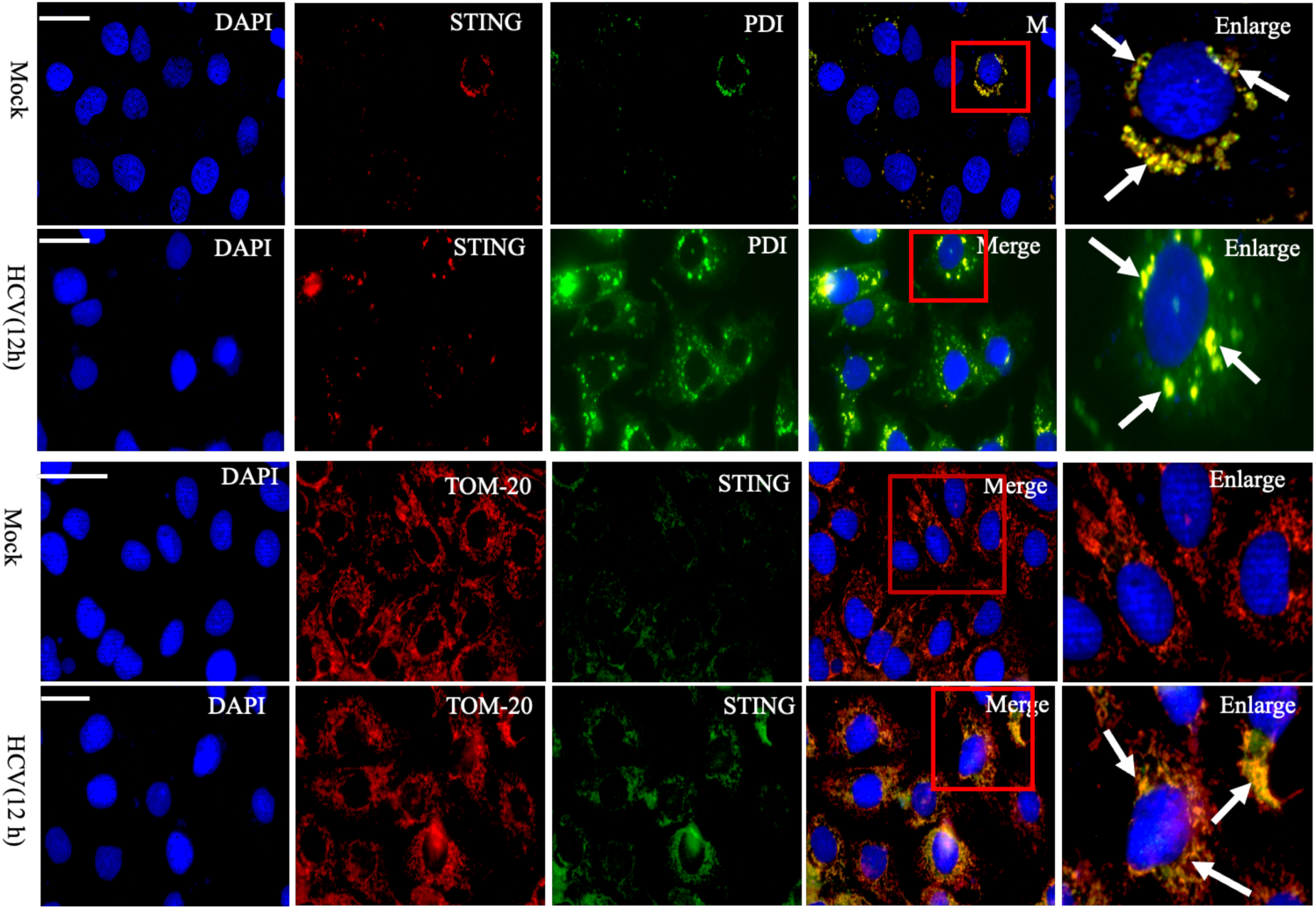
STING associates with endoplasmic reticulum and mitochondria at early infection. Huh7 (mock) and HCV infected cells were fixed, permeabilized and stained with anti-STING (mouse), anti-PDI (rabbit) for STING-ER combination, and anti-TOM-20 (mouse) and anti-STING (rabbit) primary antibodies for TOM20-STING combination. After washing with PBS, secondary antibodies Alexa Fluor-594 conjugated goat anti-mouse (red), and AlexaFluor-488 conjugated goat anti-rabbit (green) were used to study colocalization of STING with endoplasmic reticulum and mitochondria. DAPI was used to stain nuclei. Yellow dotes indicated by white arrows in the merge picture depicts the co-localization of two proteins. Images were captured at 63x magnification with a scale bar of 20μm under fluorescence microscope (Zeiss Axio observer 7).

### RIG-I, MAVS and STING co-localization dynamics during HCV infection

Viruses, including HCV, manipulate interferon (IFN) signaling to establish persistent replication within host cells. In the case of HCV, the NS4B protein directly binds to STING, an ER-resident scaffolding protein associated with MAVS (Cardif), disrupting the interaction between STING and MAVS and thereby inhibiting the antiviral signaling pathway (32). This disruption is crucial as several studies have shown that STING interacts with RIG-I and MAVS in a complex stabilized upon virus infection (33), underscoring its fundamental role in transmitting RIG-I signaling. Consequently, many RNA viruses, including HCV, have evolved strategies to block STING-dependent innate immunity. For instance, the serine protease NS4B of HCV competes with MAVS for binding to STING on mitochondria-associated membranes, displacing MAVS from the RIG-I/MAVS/STING complex and impeding downstream signaling pathways (33). In our study, we aimed to investigate whether the activation of cGAS and STING observed at 12 hours post-HCV infection plays a role in triggering the RIG-I-MAVS pathway as a host strategy to amplify the immune response against the virus. Huh7 and HCV-infected cells were fixed, permeabilized, and immunostained using antibodies against STING, RIG-I, and MAVS. Our findings showed that STING colocalizes with both RIG-I and MAVS at 12 hours post-infection, indicating that the proximity of these signaling molecules may enhance signal transduction and amplifies the antiviral response. This interaction likely facilitates more efficient production of type I interferons and other antiviral molecules, strengthening the defense mechanisms of host. RIG-I is well-known for detecting HCV dsRNA and driving MAVS-IRF3 antiviral signaling pathways (17). To demonstrate if cGAS and STING did not sense HCV dsRNA, then they may stabilize RIG-I - dsRNA binding for a stable antiviral response as knockdown of cGAS and STING have shown to upregulate the replication of many RNA viruses (34). Our immunostaining results showed significant colocalization of STING with RIG-I at 12 hours post-infection, indicating their role in the RIG-I mediated antiviral response (**Fig. 4**), which warrants further verification.

**Figure 4.**
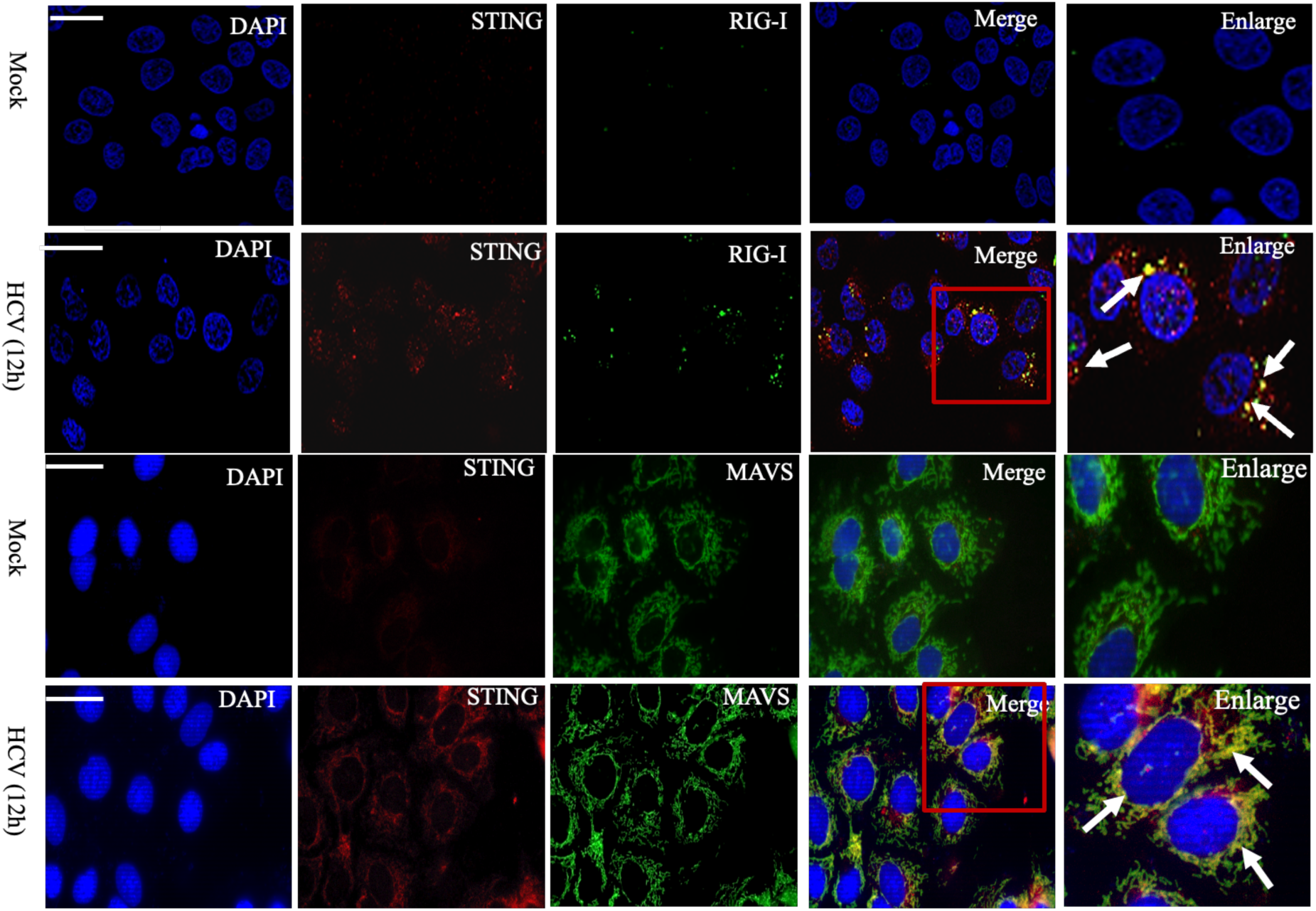
STING associates with MAVS and RIG-I at early infection. Huh (mock) and HCV infected cells were fixed, permeabilized and stained with anti-STING (mouse), anti-MAVS (rabbit) for STING-MAVS combination, and anti-STING (mouse) and anti-RIG-I (rabbit) primary antibodies for STING-RIG-I combination. After washing with PBS, secondary antibodies Alexa Fluor-594 conjugated goat anti-mouse (red), and AlexaFluor-488 conjugated goat anti-rabbit (green) were used to study colocalization of STING with MAVS and RIG-I. DAPI was used to stain nuclei. Yellow dotes indicated by white arrows in the merge picture depicts the co-localization of two proteins. Images were captured at 63x magnification with a scale bar of 20μm under fluorescence microscope (Zeiss Axio observer 7).

### cGAS and STING interaction during HCV infection

After confirming the activation of cGAS and STING at 12 hours post-HCV infection, we further examined their colocalization by immunofluorescence analysis. Mock and HCV infected (12h) Huh 7 cells were fixed and permeabilized, then primary antibodies against cGAS and STING were used followed by fluorescently labeled secondary antibodies. The merged images revealed colocalization, with the green fluorescence of cGAS and the red fluorescence of STING combining to produce a yellow signal (**Fig. 5a**). This suggests a physical and likely functional interaction between cGAS and STING during the antiviral response to HCV infection. The enhanced colocalization observed at the 12-hour time point indicates that these proteins work together in a coordinated manner, contributing to the innate immune response in the early stages of HCV infection.

**Figure 5.**
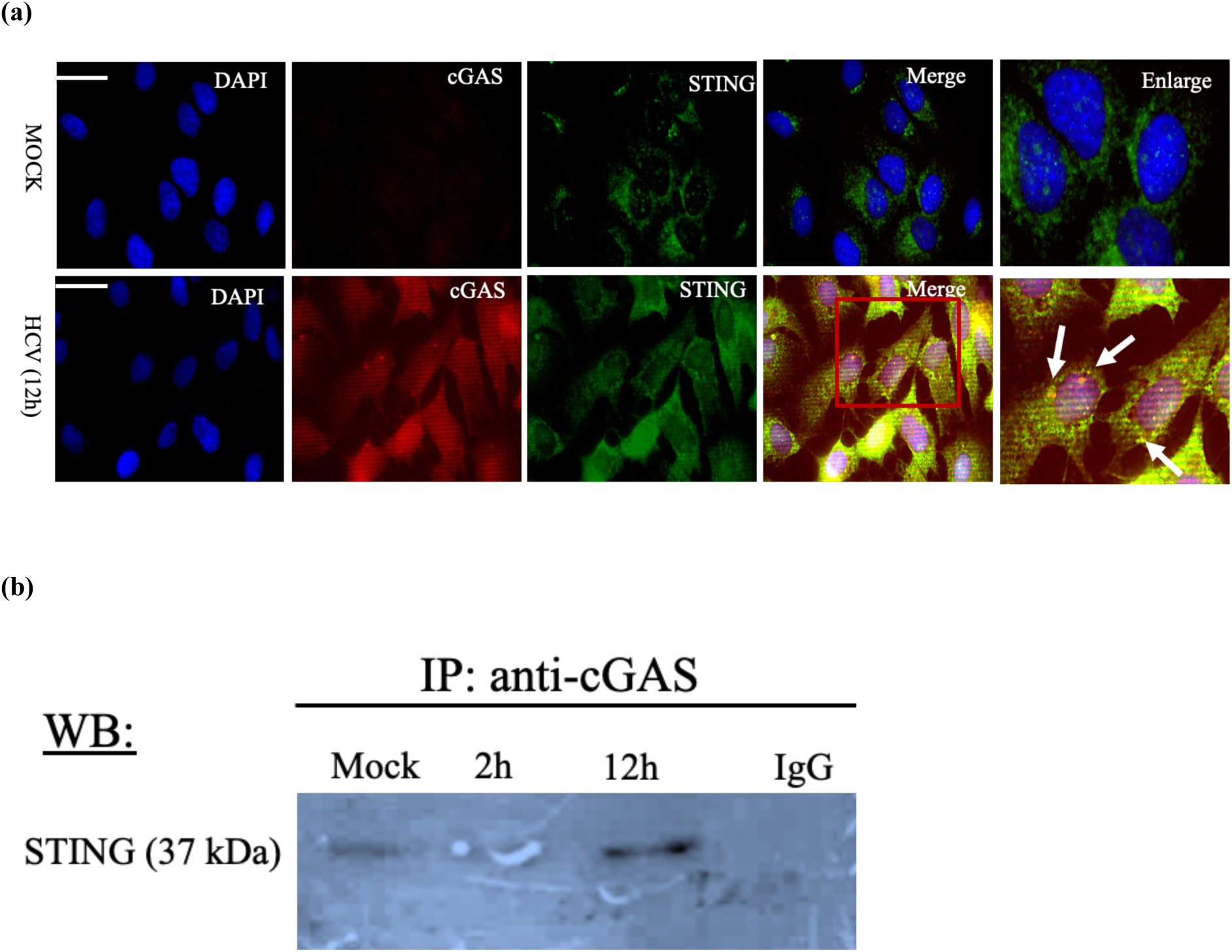
(a) cGAS associates with STING at early infection. Huh 7 (mock) and HCV infected cells were fixed, permeabilized and stained with anti-STING (mouse), anti-cGAS (rabbit) primary antibodies for STING-cGAS combination. After washing with PBS, secondary antibodies Alexa Fluor-594 conjugated goat anti-mouse (red), and AlexaFluor-488 conjugated goat anti-rabbit (green) were used to study colocalization of cGAS with STING. DAPI was used to stain nuclei. Yellow dotes indicated by white arrows in the merge picture depicts the co-localization of two proteins. Images were captured at 63x magnification with a scale bar of 20μm under fluorescence microscope (Zeiss Axio observer 7). **(b) cGAS interacts with STING at early infection:** cGAS antibody was used to pull down STING in mock and HCV-infected Huh7 cells at 2h and 12 hours post-infection. The presence of STING was confirmed by Western blot, indicating a potential interaction. IgG was used as a control antibody.

To further validate the interaction between cGAS and STING during HCV infection, Huh7 cells infected with HCV for 12 hours were lysed, and cGAS-specific antibody was used for IP. The immunoprecipitated complexes were then analyzed by Western blot using STING antibody. Our results revealed distinct STING bands in the cGAS immunoprecipitated samples at 12 hours post-infection compared to mock. In contrast IgG was used as an antibody control. However, the cGAS band could not be detected, likely due to its molecular weight being similar to the antibody heavy chain. These findings confirm the interaction between cGAS and STING, highlighting their coordinated role in the innate immune response during the early stages of HCV infection (**Fig. 5b**).

### Determining whether cGAS senses mitochondrial DNA released during early HCV infection

Mitochondrial transcription factor A (TFAM) plays a crucial role in the maintenance and regulation of mitochondrial DNA (mtDNA) (35). It plays a pivotal role in regulating the packaging, stability, and replication of the mitochondrial genome (35). Recent studies have emphasized role of TFAM in the inflammatory response triggered by mtDNA stress (36). Oxidatively damaged mitochondria can release small mtDNA fragments into the cytosol, especially when mitochondrial membrane permeability is increased (37). Dengue virus infection induces mitochondrial stress, resulting in the release of mitochondrial DNA into the cytoplasm. This DNA is detected by cGAS, which in turn activates the cGAS-STING signaling pathway (38). HCV infection induces ER stress and causes ROS production by mitochondria at the later time point of infection (39), this may also activate cGAS through the release of mtDNA which may act as PAMPs. While TFAM protects mtDNA from damage, TFAM-deficient cells release more mtDNA fragments into the cytosol, leading to higher cGAS-STING activity, evidenced by upregulated ISG expression and enhanced antiviral response (40). Moreover, proteins such as bacterial and mitochondrial nucleoid proteins HU and TFAM, along with high-mobility group box 1 protein (HMGB1), have been shown to stimulate long DNA sensed by cGAS (41). Given the observed induction of cGAS and STING at 12 hours post-HCV infection, we aimed to determine whether HCV infection triggers mitochondrial stress, leading to the release of mtDNA into the cytoplasm, where it could be sensed by cGAS and activate the cGAS-STING pathway. To investigate this, we performed colocalization studies on fixed and permeabilized Huh7 cells infected with HCV for 12 hours, followed by immunostaining for TFAM and cGAS using their respective antibodies. Our results showed expression of both cGAS and TFAM at this time point. However, no significant colocalization between these proteins was detected in the cytosolic region, suggesting that mitochondrial stress and mtDNA release do not appear to drive cGAS-STING activation at this early stage of infection (**Fig. 6**). Further investigation is needed to explore these dynamics.

**Figure 6.**
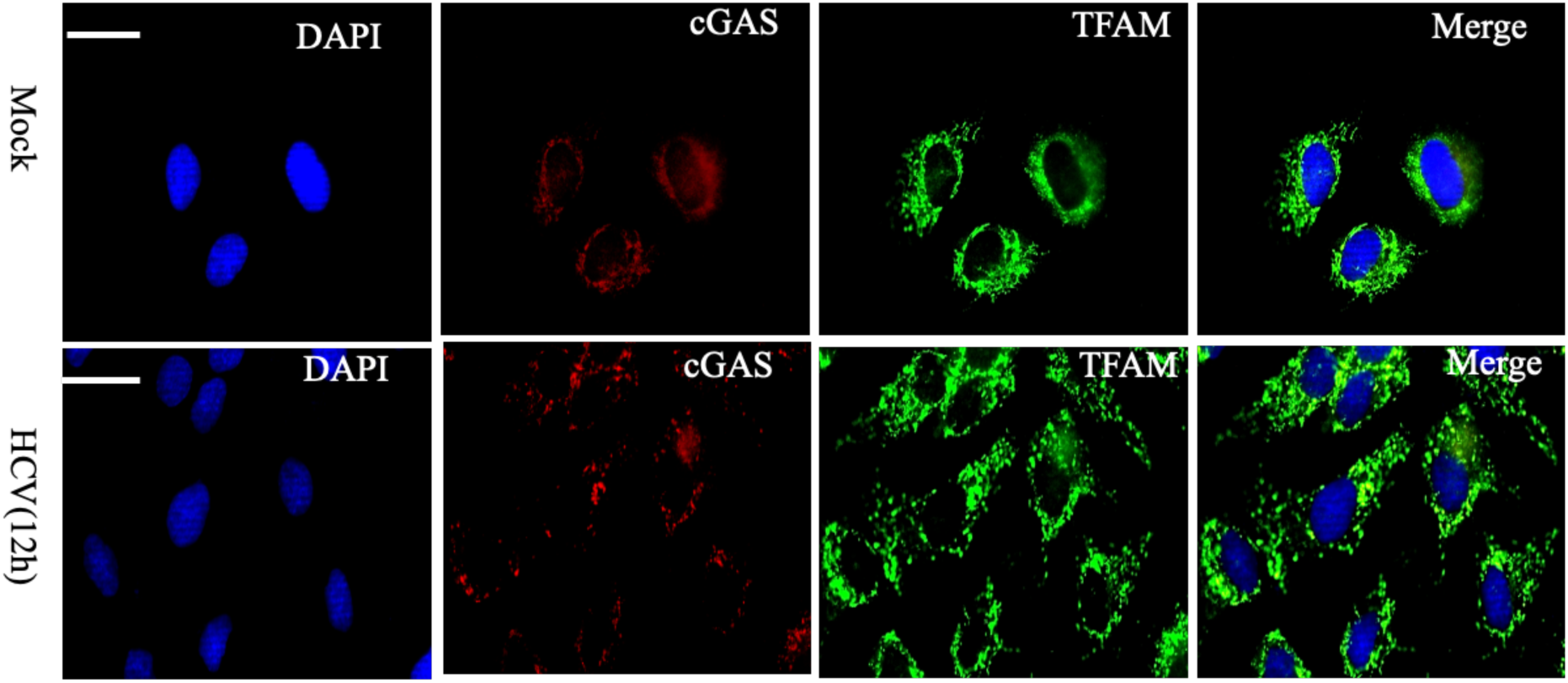
Colocalization of cGAS and TFAM at early HCV infection. Huh 7 (mock) and HCV infected cells were fixed, permeabilized and stained with anti-cGAS (mouse), anti-TFAM (rabbit) primary antibodies. After washing with PBS, secondary antibodies Alexa Fluor-594 conjugated goat anti-mouse (red), and AlexaFluor-488 conjugated goat anti-rabbit (green) were used to study colocalization of cGAS with TFAM. DAPI was used to stain nuclei. Images were captured at 63x magnification with a scale bar of 20μm under fluorescence microscope (Zeiss Axio observer 7).

### cGAS association or colocalization with HCV dsRNA during infection

Several studies have proposed that cGAS possesses RNA-binding activity, playing a key role in the cGAS-mediated innate antiviral response. This hypothesis is supported by evidence of direct RNA binding to cGAS, the identification of RNA-binding protein (RBP) partners of cGAS, and the structural resemblance between cGAS and the double-stranded RNA recognition receptor 2’-5’ oligoadenylate synthase (OAS)(42). Our findings demonstrated the activation of both cGAS and STING during HCV infection, beginning at 2 hours and peaking at 12 hours post-infection. Notably, at these time points, there was no evidence of mitochondrial stress leading to the release of mtDNA into the cytoplasm for cGAS sensing (**Fig. 6)**.cGAS is well known to requires it’s binding with ∼ 40 bp of DNA to generate cGAMP production (43). Here we used dsRNA antibody which is capable of binding ≥40 bp length of dsRNA (Cat no.# MABE1134). To verify if the activation of cGAS and STING is triggered by the sensing of dsRNA by cGAS, we conducted immunostaining using antibodies specific to cGAS and dsRNA. Our findings revealed a partial colocalization of cGAS with dsRNA at 2 hours post-infection, which became notably more pronounced at 12 hours, indicating that cGAS may directly detect HCV dsRNA during infection (**Fig. 7**). However, further validation is required to confirm this interaction. Collectively, these findings indicate that the activation of the cGAS-STING pathway is likely due to HCV dsRNA sensing by cGAS which requires further verification.

**Figure 7.**
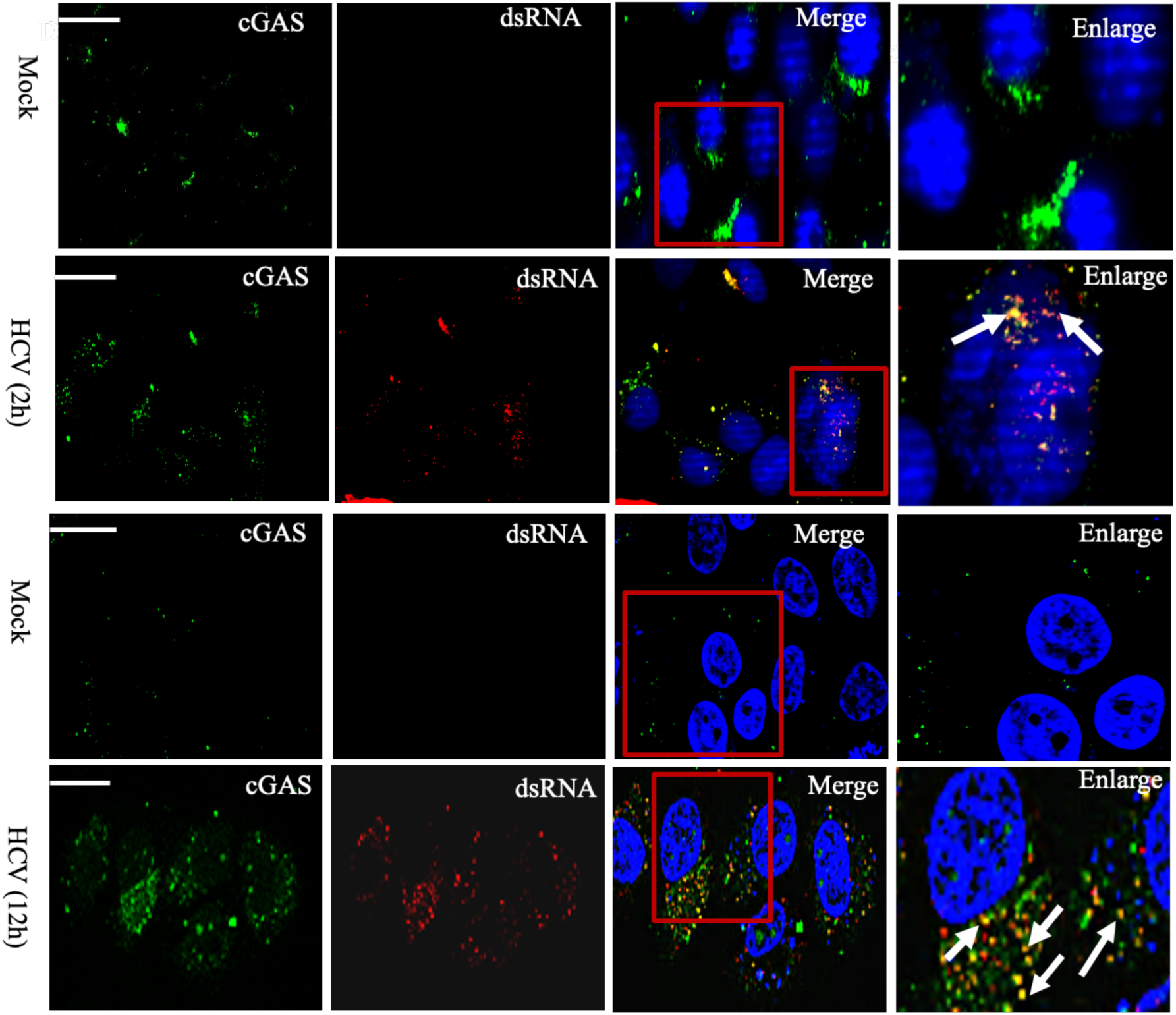
cGAS associates with dsRNA at early infection. Huh 7 (mock) and HCV infected cells were fixed, permeabilized and stained with anti-cGAS (rabbit), anti-dsRNA (mouse) primary antibodies. After washing with PBS, secondary antibodies Alexa Fluor-594 conjugated goat anti-mouse (red), and AlexaFluor-488 conjugated goat anti-rabbit (green) were used to study colocalization of cGAS with dsRNA. DAPI was used to stain nuclei. Yellow dotes indicated by white arrows in the merge picture depicts the co-localization of two proteins. Images were captured at 63x magnification with a scale bar of 20μm under fluorescence microscope (Zeiss Axio observer 7).

### Regulation of cGAS expression in STING and RIG-I depleted HCV-infected cells

The roles of cGAS and STING are well-documented in the context of many DNA viruses but only a few reports are known in RNA viruses (42). In contrast, RIG-I is known for its role as a dsRNA sensing protein (17). Our study has demonstrated an increased expression of both cGAS and STING during HCV infection **(Fig. 1a-b)**, which correlates with the subsequent activation of IFN-beta expression (**Fig. 1c**). The observed interaction between dsRNA and cGAS at an early stage of HCV infection further suggests a significant role for cGAS in antiviral response against HCV (**Fig. 7**). To confirm the involvement of cGAS and STING in the antiviral response during HCV infection, we conducted knockdown experiments targeting these genes. Huh7 cells were transfected with siRNA specific for cGAS, STING, and RIG-I. After 36 hours of siRNA treatment, cells were infected with HCV at an MOI of 1 and harvested 12 hours post-infection. Efficiency of the knockdown was confirmed, showing significant reduction in target gene expression (**Fig. 8a-c**).

**Figure 8.**
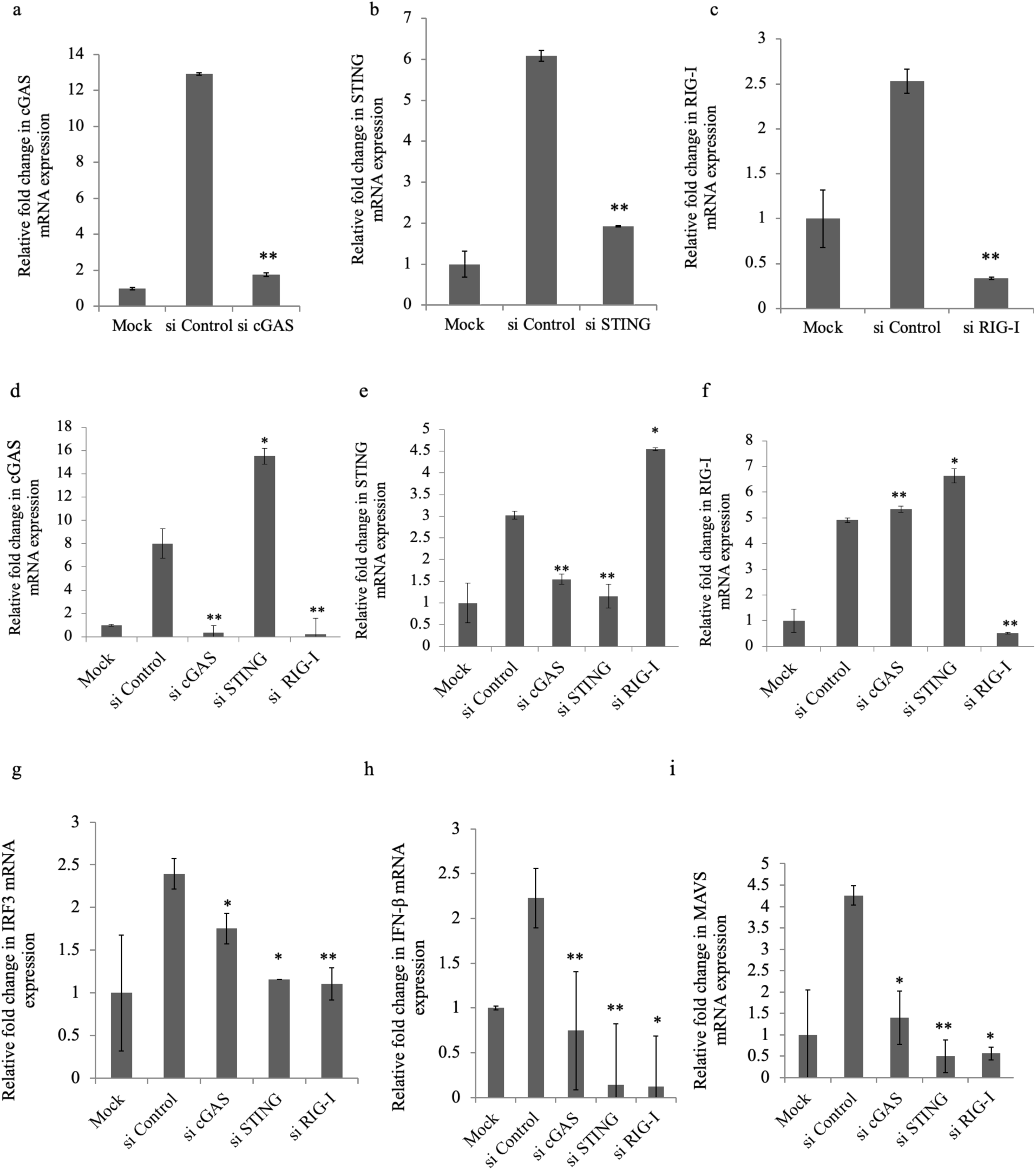
Effect of cGAS, STING, and RIG-I knockdown on the expression of cGAS, STING, RIG-I, IRF3, and IFN-β in HCV-infected Huh7 cells. Mock cells were transfected with siRNAs for 36 hours, followed by HCV infection for 12 hours, maintaining the transfection for 48 hours and infection for 12 hours. Total RNA was extracted, and RT-qPCR was performed.(a-c) Relative fold changes in cGAS, STING, and RIG-I gene expression, respectively, in cGAS, STING, and RIG-I depleted cells infected with HCV compared to the control. The values represent means ± SDs from two independent experiments performed in duplicate. (d-f) The figure shows the relative fold change in cGAS expression in STING and RIG-I depleted cells compared to control. The figure shows the relative fold change in STING gene expression in cGAS and RIG-I depleted cells compared to control. The figure shows the relative fold change in RIG-I expression in cGAS and STING depleted cells compared to control (g-i) effect of cGAS, STING, and RIG-I knockdown on IRF3, IFN-β and MAVS gene expression. The figure shows the relative fold change in IRF3 and IFN-β expression in cGAS, STING, and RIG-I depleted cells compared to control. *p-value ≤ 0.05, **p-value ≤ 0.01 for si Control vs si cGAS, si STING, and si RIG-I.

First, we examined cGAS induction in siControl cells (HCV infected Huh 7 cells at 12h post infection) and observed a significant increase in cGAS levels compared to Mock cells. Next, we checked the cGAS expression in STING and RIG-I knockdown (KD) cells compared to control siRNA (siControl). We found an increased expression of cGAS in STING KD cells. This result suggest that STING suppresses cGAS expression that could be possible when HCV NS4B targets STING, it may exploit STING to suppress cGAS to abolish cGAS-STING pathway. (**Fig. 8d**). Conversely, knockdown of RIG-I led to decreased expression of cGAS, highlighting that RIG-I may play a supportive role in maintaining cGAS expression and its associated signaling pathways during HCV infection (**Fig. 8d**). Since, HCV NS3/4A cleaves MAVS and disrupt RIG-I-MAVS antiviral pathway, this could be a novel role of RIG-I alone to stabilize or enhance cGAS to maintain antiviral state through cGAS-STING pathway.

### Modulation of STING expression in absence of cGAS and RIG-I during HCV infection

Detection of viral genome by cGAS can lead to the production of cGAMP, a crucial molecule for STING activation (44). To investigate the impact of cGAS knockdown on STING expression during HCV infection and to determine whether this pathway is active, we performed targeted gene knockdown experiments. Huh7 cells were transfected with siRNA specific to cGAS, STING, and RIG-I (36 h), followed by HCV infection for 12 hours. RNA was isolated from these cells, and RT-qPCR was employed to quantify the mRNA expression levels of the target genes. Our results revealed STING expression compared to mock. However, knockdown of cGAS significantly reduced the mRNA expression of STING (**Fig. 8e**). This indicates that cGAS is a critical upstream regulator necessary for cGAMP production followed by STING activation during HCV infection. Furthermore, we observed an increased expression of STING in RIG-I knockdown cells (**Fig. 8e**), suggesting that RIG-I limit STING transcript during HCV infection which requires more study to uncover the mechanism. These findings suggests the complex regulatory network among cGAS, STING, and RIG-I in the innate immune response against HCV.

### Effect of cGAS and STING on RIG-I expression during HCV infection

After transfecting Huh7 cells with siRNA targeting cGAS and STING and RIG-I, followed by HCV infection for 12 hours, we analyzed the mRNA expression levels using RT-qPCR. Our results showed a significant increase in RIG-I expression in control siRNA compared to (mock) cells (**Fig. 8f**). However, we observed some increase in RIG-I expression in STING knockdown cells, but some effect was also observed in cGAS knockdown cells (**Fig. 8f**). The upregulation of RIG-I in absence of STING suggests that STING limits RIG-I expression (**Fig. 8f**). Likewise increase in STING expression in absence of RIG-I also indicated that RIG-I limits STING expression (**Fig. 8e**). These findings suggesting that STING and RIG-I are antagonistic to each other. Since, RIG-I-MAVS and cGAS-STING pathways are generated by host in response to viral infection, it could be viral strategy to abolish cGAS-STING pathway as it does with RIG-I-MAVS using NS3/4A. It seems virus is targeting host proteins like STING or MAVS which needs further study. The colocalization of STING with RIG-I in our immunofluorescence assays (**Fig. 4**) further strengthen that these molecules are associating together in a coordinated manner to amplify the antiviral response at early stages of HCV infection. However, these proteins are also target to each other to limit the antiviral response which could be viral strategy.

### Indispensable role of of cGAS, STING and RIG-I on IRF3 expression during HCV infection

Next, we sought to determine the depletion effect of cGAS, STING, and RIG-I on downstream gene expression. Our primary focus was on IRF3, a key factor in activating the interferon response once the cGAS-STING and RIG-I-MAVS pathway is activated (23). We observed that IRF3 expression was increased in HCV infected cells compared to mock, which was significantly decreased in cGAS and STING depleted cells **(Fig. 8g)**. Additionally, IRF3 expression was reduced in RIG-I-depleted cells (**Fig. 8g**). These findings suggests the critical importance of the cGAS-STING pathway during HCV infection apart from RIG-I-MAVS pathway. The knockdown of cGAS leading to diminished IRF3 expression indicates that cGAS is crucial for the upstream signaling necessary for IRF3 activation. Similarly, the reduction of IRF3 expression following STING knockdown highlights the essential role of the cGAS-STING pathway in mediating IRF3 activation. The decreased IRF3 expression in RIG-I-depleted cells suggests that RIG-I also plays a role in the IRF3-mediated interferon response at early infection which is abolished at later infection when NS3/4A cleaves MAVS as shown earlier (19). Collectively, these results demonstrate that cGAS, STING, and RIG-I are essential for the proper activation of IRF3, a key transcription factor in the antiviral immune response against HCV infection, also these findings indicate that both pathways are interconnected to each other which requires further investigation.

### cGAS, STING and RIG-I enhance IFN-β induction during HCV infection

To further confirm the importance of the cGAS-STING pathway during HCV infection, we assessed the status of IFN-β, a key molecule in the antiviral response, in cGAS and STING knockdown cells. Our results showed a significant increase in IFN-β expression in control siRNA compared to (mock) cells. We observed a significant decrease in IFN-β expression in cGAS and STING knockdown cells (**Fig. 8h**). This suggests the critical role of the cGAS-STING pathway in mounting an effective immune response against HCV. The concurrent reduction in IRF3 and IFN-β levels confirms the activation of the cGAS-STING pathway during HCV infection. Additionally, depletion of IFN-β in RIG-I knockdown cells (**Fig. 8h**), indicates the importance of established RIG-I -MAVS in the antiviral response to HCV infection. These findings highlight the essential function of the cGAS-STING-RIG-I axis in inducing IFN-β and suggest that these proteins work together to elicit a robust antiviral response against HCV. The observed downregulation of IFN-β upon knockdown of these genes validates their critical role in the innate immune response to HCV infection.

### cGAS, STING and RIG-I induces MAVS expression during HCV infection

Mitochondrial antiviral-signaling protein (MAVS) is a vital component of the innate immune response against viral infections, including hepatitis C virus (HCV) (17),(19). Located on the outer membrane of mitochondria, MAVS is central to the antiviral signaling pathway activated by the recognition of viral RNA through RIG-I-like receptors (RLRs) (17). Our findings show that MAVS colocalizes with STING **(Fig. 4)** prompting us to examine the impact of cGAS and STING depletion on MAVS. We observed increased MAVS expression in HCV infected cells transfected with Sicontrol compared to mock. However, decrease in MAVS expression was observed in cGAS and STING knockdown cells (**Fig. 8i**). Further, the MAVS expression was decreased in RIG-I depleted cells which is expected based on previous studies (45).Together, this reduction highlights the critical role of cGAS-STING pathway in maintaining MAVS levels during HCV infection. Additionally, in the absence of RIG-I, the cGAS-STING pathway may compensate by forming a complex with MAVS (STING-MAVS, STING-RIG-I) to activate the innate immune response, suggesting the importance of both pathways during HCV infection.

### cGAS and STING modulates HCV proteins during infection

Studies have shown that HCV NS3 and HCV NS5A are crucial for its replication, however HCV NS5A and Core are widely studied for their role in HCV virion assembly. Based on the previous findings, here we have determined the expression of these viral proteins in knockdown studies.

To investigate the roles of cGAS and STING during HCV infection, we examined their knockdown effect on HCV proteins which are essential for replication and assembly. Our results showed a significant increase in HCV NS3, core and NS5A expression in control siRNA compared to (mock) cells. cGAS depletion resulted in increased expression of both HCV core and NS5A proteins, while having not much effect on NS3 level (**Fig. 9a-c**). This suggests that cGAS negatively regulates the expression of core and NS5A, possibly by activating antiviral responses that suppress viral protein synthesis. Conversely, STING depletion led to increased expression of NS3 protein, indicating a potential inhibitory role of STING on NS3 levels, but it showed decrease in NS5A mRNA expression and some increase in core mRNA expression (**Fig. 9a-c**). These findings indicate the distinct roles of cGAS and STING in modulating HCV protein expression, suggesting that cGAS and STING broadly impact viral proteins regulation, reflecting their different functions in the innate immune response against HCV infection.

**Figure 9.**
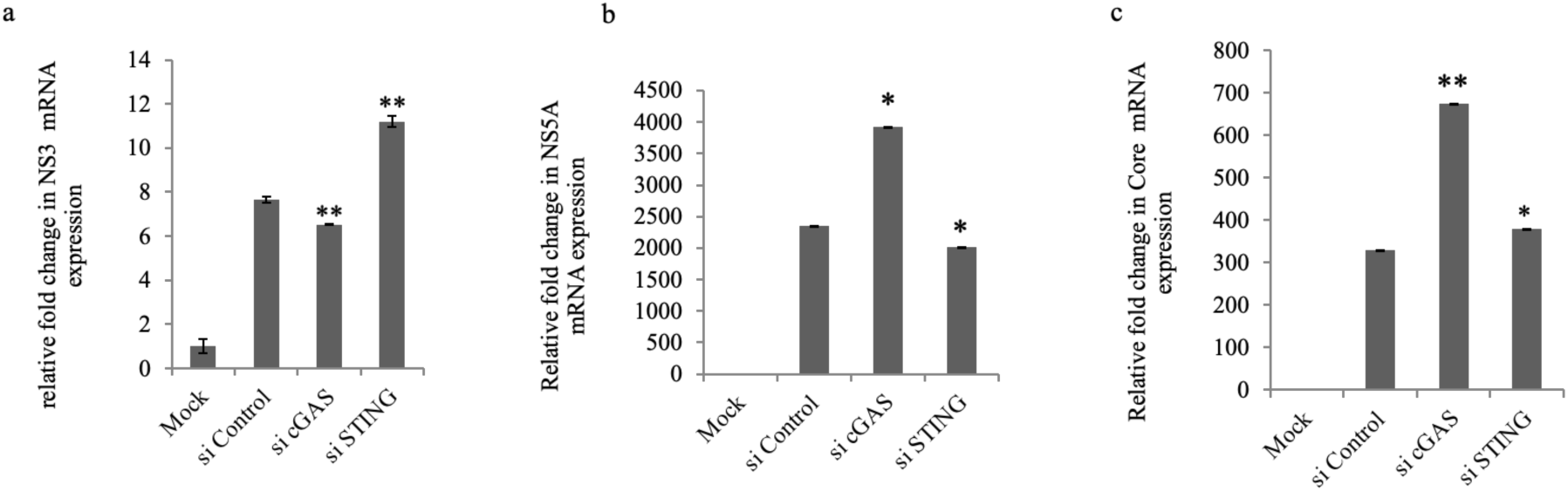
cGAS and STING depletion effect on HCV proteins. Mock cells were transfected with siRNAs for 36 hr followed by HCV infection for 12h to maintain the transfection for 48 hr and infection for 12h. Total RNA was extracted and RT-qPCR was performed. (a-c) Shows the relative fold change in HCV Core, NS3 and NS5A gene expression in cGAS and STING depleted cells infected for a period of 12hrs compared to control. The values represent the means ± SDs from two independent experiments performed in duplicate. * p-value ≤ 0.05, ** p-value ≤0.01, for si Control vs si cGAS and si STING.

### 2’3’-cGAMP production during HCV infection

Upon sensing the viral genome, cGAS catalyzes the formation of 2’3’-cGAMP, which then binds and activates STING to further stimulate the INF-β response during viral infection. This pathway is well-known for its role when cGAS detects the dsDNA of viruses (17). Given our findings of cGAS and STING activation during HCV infection and the observation that cGAS binds to dsRNA, we aimed to determine if cGAMP is also produced during HCV infection which is required for STING activation followed by downstream process. Huh7 cells were infected with HCV at various time points, and cell lysates were collected. cGAMP levels were quantified using a cGAMP detection kit (Cayman Chemical). We observed an increase in cGAMP production at 2 hours, on peak at 12 hours post-HCV infection (data not shown). This peak coincided with higher cGAS expression at 12 hours, suggesting activation of the cGAS-STING pathway through cGAMP during HCV infection. In contrast, we could not observe significant cGAMP production in the supernatant collected from mock and 12 hour post HCV infection. Moreover, THP1cells that were kept as positive control showed significant cGAMP production.

To further validate our results, we knocked down cGAS, STING, and RIG-I using gene-specific siRNA, along with control siRNA. Huh7 cells were transfected with the siRNAs, and 36 hours post-transfection, cells were infected with HCV for 12 hours to maintain transcription for 48 h and HCV infection for 12 h. Cell lysates were prepared, and cGAMP levels were measured using the cGAMP detection kit, following the manufacturer’s protocol. We observed increased cGAMP level in sicontrol compared to mock sample, however, a decreased cGAMP levels in cGAS knockdown cells. Similar result was observed in STING depleted cells. Interestingly, RIG-I depletion led to reduced cGAMP levels as well, which is quite expected as we observed that RIG-I enhance cGAS activation. The decrease in cGAMP in STING knockdown cells further validates the essential role of the cGAS in STING activation through cGAMP (**Fig. 10**)

**Figure 10.**
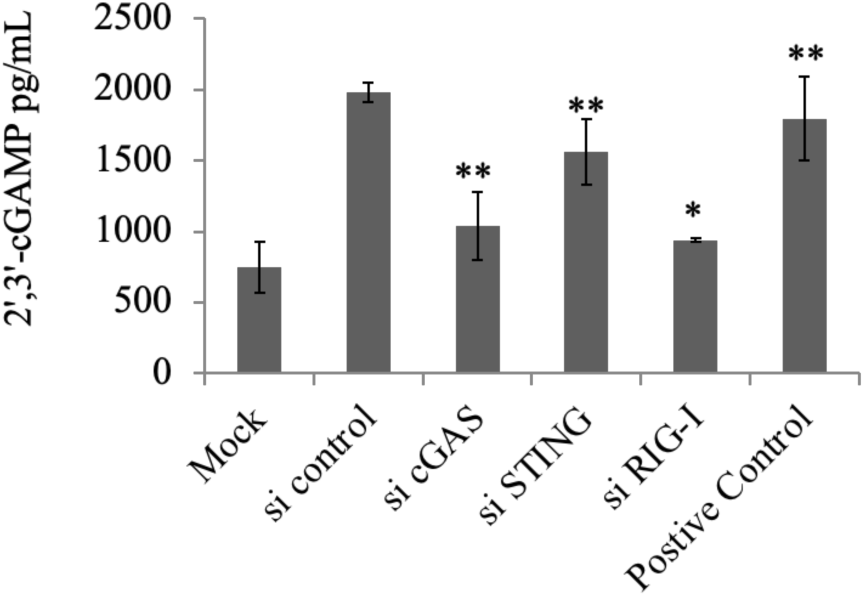
cGAS activation causes cGAMP production in early HCV infection. Huh7 (Mock) cells were transfected with siRNAs for 36 hr followed by HCV infection for 12h to maintain the transfection for 48 hr and infection for 12h. cGAS, STING and RIG-I depletion effect on cGAMP production in Huh 7 cells at 12 h of HCV infection. The values represent the means ± SDs from two independent experiments performed in duplicate. * p-value ≤ 0.05, **p-value ≤0.01, for mock vs infection and si Control vs si cGAS, si STING and si RIG-I.

## Discussion

HCV poses a significant public health challenge, leading to severe liver diseases such as cirrhosis and hepatocellular carcinoma (46). The virus is characterized by a high mutation rate, which enables it to continuously evolve (47). In India, HCV significantly contributes to the annual death toll, underscoring its impact on public health. The rapid advancements in HCV research have been greatly facilitated by the development of robust cell culture systems and small animal models, which have been instrumental in understanding the life cycle and pathogenesis of HCV (48). In our study, we utilized the JFH-1 strain of HCV genotype 2a, known for its unique ability to propagate efficiently in cell culture systems without requiring culture-adaptive mutations (49). This strain has provided invaluable insights into viral replication processes, host-virus interactions, and the efficacy of potential antiviral treatments. By leveraging the JFH-1 strain, we conducted comprehensive analyses and experiments that would have not been feasible with other HCV strains that do not propagate as effectively *in vitro*. This approach has been pivotal in advancing our understanding of HCV biology and in developing therapeutic strategies to combat the persistent virus. After viral infection the primary responsibility of the host is to detect the viral presence and strengthen the immune response. Host cells possess various nucleic acid sensors capable of distinguishing between host and viral nucleic acids. Upon sensing danger signals in the cytoplasm, these sensors initiate an innate immune response. Traditionally, cellular antiviral mechanisms in the context of HCV infection have focused on RNA sensors like RIG-I and MDA5 (17). RIG-I senses dsRNA and activates the adapter protein MAVS, triggering downstream IRF3 signaling and thereby enhancing IFN-I production (33). This pathway has been shown to be abrogated at later stages of infection when MAVS is cleaved by the HCV protease NS3/4A, resulting in the suppression of interferon production and increased viral replication (19). However, viral replication has been shown to be suppressed even in the absence of this pathway, indicating that the host has alternative strategies to reduce viral infection. Recently, it has been shown that the host protein cGAS plays an important role during RNA viral infections (50). cGAS is primarily a viral dsDNA sensing protein that detects in the cytoplasm and catalyzes the production of cGAMP (23), which is necessary to activate the signaling pathway adapter protein STING. Once STING is activated, it further activates downstream molecules, ultimately leading to the production of interferons (51). Although cGAS primarily detects cytoplasmic DNA, it has also been shown to bind RNA and defend against RNA viruses such as West Nile virus (WNV), dengue virus (DENV), and murine norovirus (MNV) (42). Numerous studies have suggested the potential RNA-binding activity of cGAS and its role in the cGAS-mediated innate antiviral response, supported by the direct binding of RNA to cGAS, the RNA-binding protein (RBP) partners of cGAS, and the structural similarity of cGAS to the dsRNA recognition receptor 2’-5’ oligoadenylate synthase (OAS) (42). In the context of HCV, it has been shown that the viral protein NS4B targets STING and disrupts the RIG-I-MAVS-STING complex, which plays a crucial role in the antiviral response (32). The question then arises: how do cGAS and STING function during HCV infection, and what is the molecular mechanism behind their activation? This study focuses on the role of cGAS and STING during HCV infection, aiming to elucidate the molecular mechanisms involved in their activation and contribution to the immune response against HCV.

The initial step of our study was to assess the status of cGAS expression, which we accomplished through a kinetic study. We specifically selected the early time point of HCV infection, as this is the time when viral genome sensing and the immune response are most active. Our investigation into the induction of cGAS revealed a marked increase in its expression at early stages of HCV infection, specifically, 12 hours, and decreases at day 1 (D1) post-infection (**Fig. 1a**). This temporal pattern of cGAS expression indicates its potential role in the initial antiviral defense. The early induction suggests that cGAS may recognize viral components soon after infection, thereby initiating downstream signaling pathways essential for mounting an effective immune response. The concomitant induction of STING and IFN-β at early time points aligns with the known cGAS-STING-IRF3 signaling axis (**Fig. 1b, c)**. The activation of STING likely facilitates the recruitment of key signaling molecules such as TBK1 and IRF3, culminating in the production of IFN-β and the establishment of an antiviral state. However, we could not detect IFN-α and IFN-γ that could be due to their minimal expression because of the nature of epithelial origin of Huh7 cells (**Supplementary** Fig. 2).

Next, the immunofluorescence studies demonstrating association of cGAS with both the endoplasmic reticulum (ER) and mitochondria during early HCV infection (**Fig. 2**), provide new insights into the spatial dynamics of antiviral signaling, where it may potentially influencing mitochondrial antiviral signaling (MAVS) pathways. This dual localization may enhance the ability of host to detect and respond to viral infections. Parallel to cGAS, STING also showed significant colocalization with ER and mitochondria in HCV-infected cells at 12 h of infection (**Fig. 3**). Interestingly we observed cGAS-STING colocalization which were verified by immunoprecipation **(Fig 5)**. Our results suggest that during HCV infection, cGAS might also be involved in sensing RNA or RNA-induced stress responses at these organelles. This expands the functional repertoire of cGAS and highlights its versatility in antiviral defense mechanisms. The observed colocalization of RIG-I-STING, and MAVS-STING at 12 hours post-infection (**Fig. 4)**, suggests the integrated nature of antiviral signaling networks. RIG-I, a well-established RNA sensor (17), appears to work in concert with STING and MAVS, forming a complex that amplifies the antiviral response. This interaction likely enhances the stability and efficiency of signal transduction, ensuring a robust production of type I interferons and other antiviral molecules. Dengue viral infection has been shown to cause mitochondrial stress, leading to the leakage of mitochondrial DNA(mtDNA) into the cytoplasm (38). This DNA is then sensed by cGAS, which activates the cGAS-STING pathway. To confirm this hypothesis in HCV infection, we conducted colocalization studies to examine the interaction between cGAS and mitochondrial DNA. Despite the significant expression of cGAS and TFAM at 12 hours post-infection, we did not observe direct colocalization **(Fig. 6)**, indicating that mitochondrial stress and mtDNA release might not be primary drivers of cGAS activation at this stage. Instead, our findings suggest that cGAS activation may be mediated through direct recognition of viral RNA or RNA-induced cellular changes. The immunostaining results showing cGAS association with dsRNA at early infection provide compelling evidence for its role in sensing viral RNA (**Fig. 7**). This challenges the traditional paradigm of cGAS as exclusively a DNA sensor and opens new avenues for understanding its function in RNA virus infections like HCV. The interaction with dsRNA suggests that cGAS might contribute to antiviral responses through mechanisms previously unrecognized which needs further verification. There is a possibility that cGAS if it is not sensing viral RNA directly then it might be assisting RIG-I-dsRNA binding for stable antiviral response.

Moreover, the knockdown experiments targeting cGAS, STING, and RIG-I in HCV-infected Huh7 cells revealed significant insights into the interplay between these molecules in the antiviral response, knockdown of STING resulted in increased expression of cGAS, suggesting a limiting effect of STING that could be due to targeting by HCV NS4B **(Fig. 8d)**. Conversely, knockdown of RIG-I led to decreased expression of cGAS, highlighting that RIG-I may play a supportive role in maintaining cGAS expression and its associated signaling pathways during HCV infection **(Fig. 8d)**. cGAS plays a pivotal role in activating STING by producing cGAMP upon viral RNA detection (23). This study demonstrated that cGAS knockdown significantly reduced STING mRNA levels, suggesting role of cGAS as an upstream regulator of STING activation during HCV infection (**Fig. 8e**). Conversely, RIG-I knockdown resulted in increased STING expression, indicating that RIG-I may limit STING transcript levels during HCV infection (**Fig. 8e**). Similarly, the upregulation of RIG-I in the absence of STING implies that STING may limit RIG-I expression (**Fig. 8f**). These findings indicate that STING and RIG-I act as antagonists to each other. Since the host generates RIG-I-MAVS and cGAS-STING pathways in response to viral infections, it is possible that the virus employs strategies to abolish the cGAS-STING pathway, as it does with the RIG-I-MAVS pathway using NS3/4A (19). This interplay between different PRRs indicating the redundancy and robustness of the innate immune system in mounting an effective defense against HCV. IRF3 is a crucial transcription factor activated by the cGAS-STING pathway, leading to the production of interferons (17). This study showed that IRF3 expression was significantly reduced in cGAS, STING, and RIG-I knockdown cells, highlighting the importance of these molecules in activating IRF3 and initiating an effective antiviral response (**Fig. 8g**). The diminished IRF3 levels upon knockdown of these genes indicate their critical role in the upstream signaling necessary for IRF3 activation. Interferon-beta (IFN-β) is a key cytokine in the antiviral response. This study observed a significant decrease in IFN-β expression in cGAS and STING knockdown cells, which was further reduced upon RIG-I depletion (**Fig. 8h**). These findings confirm the essential role of the cGAS-STING-RIG-I axis in inducing IFN-β and suggest a cooperative interaction among these components to elicit a robust antiviral response against HCV. Mitochondrial antiviral-signaling protein (MAVS) is a critical component of the innate immune response (52). This study found that MAVS expression decreased in cGAS and STING knockdown cells, indicating their role in maintaining MAVS levels during HCV infection (**Fig. 8i**). The reduction of MAVS in RIG-I depleted cells highlights the importance of RIG-I in MAVS activation. The study further investigated the roles of cGAS and STING in regulating HCV proteins. Our finding indicates that cGAS negatively regulates HCV core and NS5A expression by activating antiviral responses that suppress viral protein synthesis (**Fig. 9a-c**). Also, STING specifically targets NS3 expression, indicating its specific regulatory influence on NS3 levels. These findings highlight the distinct roles of cGAS and STING in modulating HCV protein expression and maintain antiviral response at early infection.

The formation of 2’3’-cGAMP by cGAS upon sensing the viral genome is critical for activating STING and initiating the INF-β response during viral infection. This pathway is well-established for cGAS detection of dsDNA from viruses. Our results demonstrate that cGAS and STING are activated during HCV infection, with cGAS binding to dsRNA (**Fig. 7**) indicating its role in HCV infection. The reduction in cGAMP levels in cGAS knockdown cells (**Fig. 10**) confirms the direct involvement of cGAS in detecting HCV and activating downstream signaling. Additionally, decreased cGAMP levels in STING knockdown cells indicating the critical role of STING in this pathway, reinforcing the essential nature of the cGAS-STING axis in the innate immune response. Furthermore, the reduction in cGAMP levels in RIG-I knockdown cells highlights crucial role of RIG-I in maintaining cGAS induction (**Fig. 10**). This suggests that RIG-I may influence cGAS activation or that these pathways work together as part of a coordinated antiviral response. This interaction emphasizes the complexity and redundancy of the host antiviral defense mechanisms, ensuring robust protection against HCV infection. Our findings provide a deeper understanding of the molecular mechanisms involved in HCV sensing and the innate immune response, potentially guiding the development of new therapeutic strategies targeting the cGAS-STING pathway for treating HCV and other viral infections.

## Materials and Methods

### Reagents and chemicals

Dulbecco’s Modified Eagle Medium (DMEM), Fetal Bovine Serum (FBS), Phosphate-Buffered Saline (PBS), 4-(2-Hydroxyethyl)-1-piperazineethanesulfonic Acid (HEPES, 1M), L-Glutamine (100X), Non-Essential Amino Acids (100X), Antibiotics (Penicillin, Streptomycin, Amphotericin B, 100X), and TriZol were all sourced from Thermo Fisher Scientific (GibcoTM, Waltham, MA, USA). Cell culture dishes were obtained from Corning, and microscopic glass cover slips from Blue Star. Molecular grade paraformaldehyde and Triton X-100 were purchased from Thermo Fisher Scientific, along with the cDNA synthesis kit. The KAPA SYBR FAST qPCR Master Mix (2X) and molecular grade ethanol were purchased from Merck. Thermo Fisher Scientific supplied Tris base, NaCl, glycine, APS, PBS, and Bovine Serum Albumin (BSA). BIO-RAD (India) provided the 30% acrylamide and bis-acrylamide solution, while the PVDF membrane was sourced from PALL Life Sciences (USA). X-Ray films were acquired from Kodak. The 2’3’-cGAMP ELISA Kit (501700) was purchased from Cayman Chemical (USA), and DAPI and molecular grade DMSO were sourced from Merck (India).

### Antibodies utilized in this study

This study utilized various antibodies for Western blotting and immunofluorescence. The following table details each antibody used in this study.

| Antibody name | Source | Purpose | Purchased from |
| --- | --- | --- | --- |
| STING | Anti-Mouse and Anti-Rabbit | WB/IFA | BIO-RAD (VMA00456) and CST (D2P2F) |
| cGAS | Anti-Rabbit and Anti mouse | WB/IFA | CST (E5V3W), Proteintech (264116-1-AP) and MyBioSource (MBS9217055) |
| MAVS | Anti-Rabbit | WB/IFA | BIO-RAD (AHP1702) |
| Actin | Anti-Rabbit | WB/IFA | Cusabio technology LLC |
| Tom20 | Anti-Rabbit | WB/IFA | Cell Signaling Technology |
| HCV core | Anti-mouse | WB/IFA | GeneTex International corporation (GTX41719) |
| RIG-I | Anti-Rabbit | WB/IFA | Cell Signaling Technology (D33H10) |
| Ant-dsRNA | Anti-Mouse | IFA | Millipore-Sigma (MABE1134) |
| TFAM | Anti-Mouse | IFA | Thermofisher (18G102B2E11) |
| PDI | Anti-Rabbit | IFA | Cell signaling technology (C81H6) |

### List of Primers

All the primers used in the study were purchased from Bioserve Biotechnologies (India) and are listed in the given table

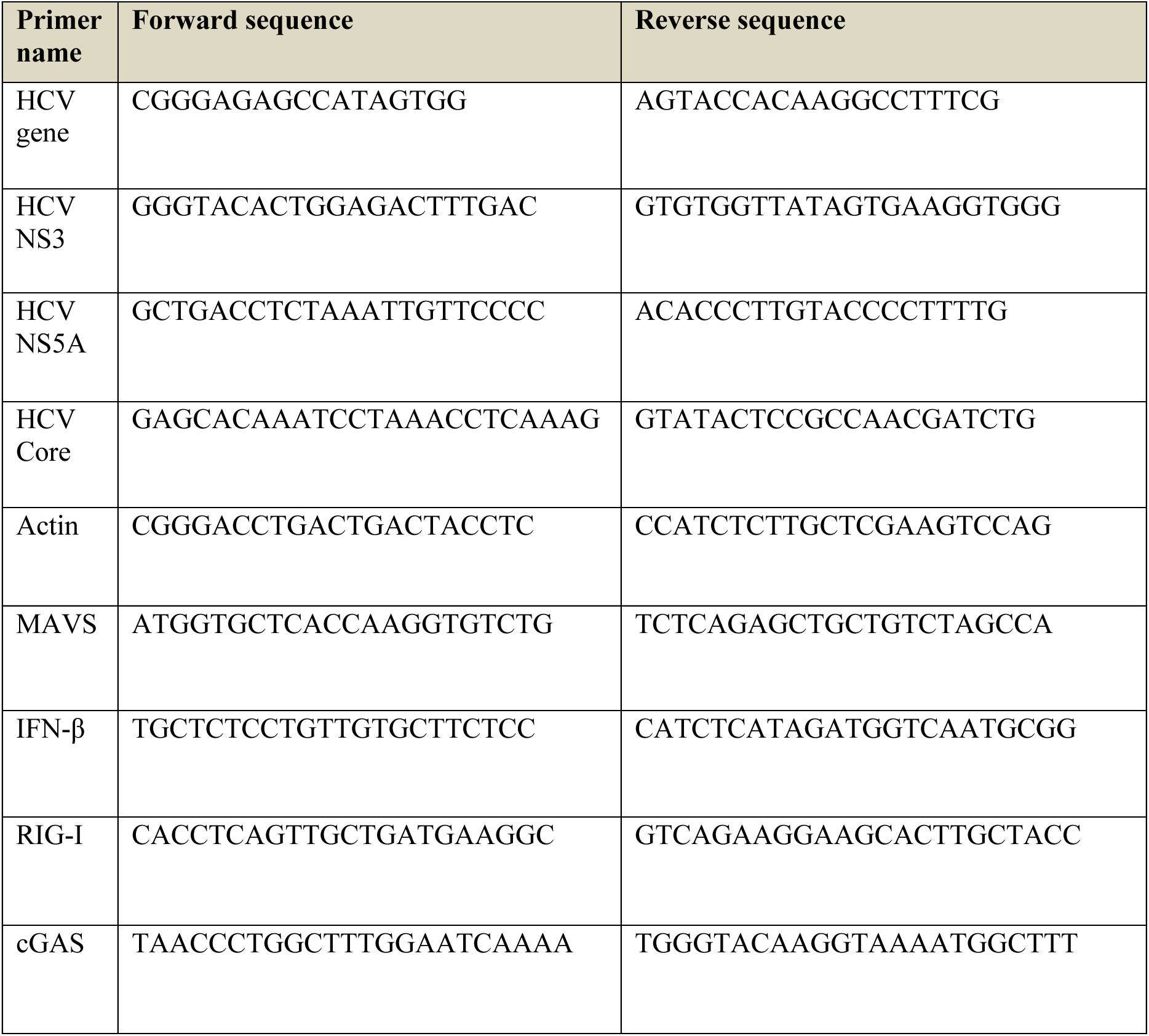
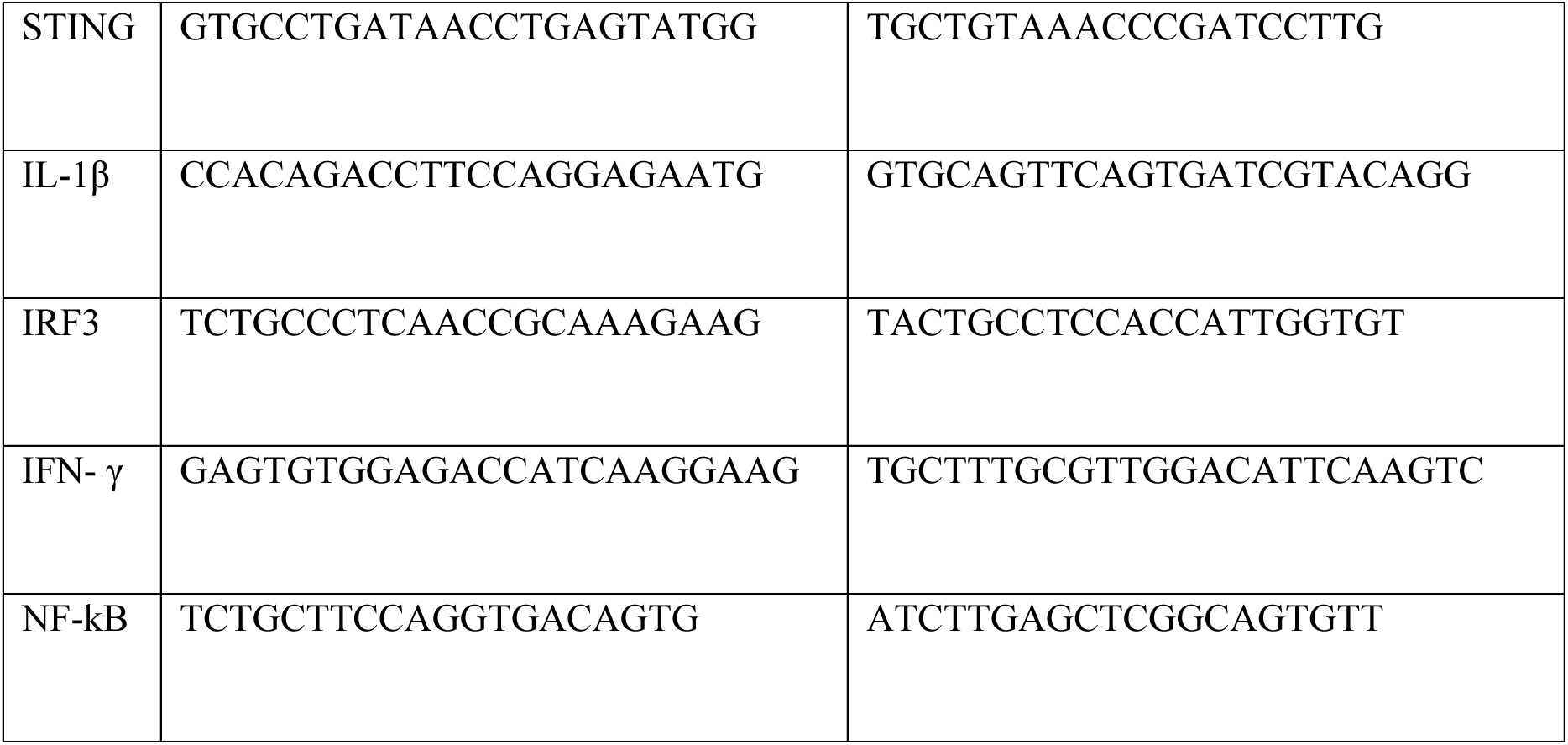

### Cell lines and HCV cell culture system

Human hepatoma cell line, Huh7 was obtained from Dr. Ranjith Kumar at the University School of Biotechnology, Guru Gobind Singh Indraprastha University, New Delhi, India. The Huh7 cells were cultured in Dulbecco’s Modified Eagle Medium (DMEM) provided with 10% fetal bovine serum (FBS), 1% penicillin-streptomycin antibiotic cocktail, 1% HEPES buffer, and 1% non-essential amino acids. The cells were maintained in a humidified incubator with 5% CO2 at 37°C. Cells were passaged when they reached 80-90% confluence, and the media was refreshed every 24-48 hours to ensure optimal growth conditions. Plasmids for HCV genotype 2a, including J6/JFH-1 and the replication-defective JFH-1/GND, were obtained from Apath L.L.C., Rockefeller, USA under a material transfer agreement. HCV particles were generated following established protocols (53).

### RNA isolation from cells using TRIzol method

RNA was isolated from the cells using a modified TRIzol method (54). Briefly, the media was removed from the cells, and 1 mL of TRIzol was added to an 80% confluent 60 mm dish. The cells were lysed thoroughly by pipetting up and down. After adding 200 μL of chloroform, the mixture was vortexed for 15 seconds and left to stand at room temperature for 10 minutes. The sample was then centrifuged at 12,000 rpm for 15 minutes at 4°C to achieve clear phase separation. The aqueous phase was carefully transferred to a fresh microcentrifuge tube, and 500 μL of isopropyl alcohol was added to precipitate the RNA. The solution was incubated at room temperature for 30 minutes and then centrifuged at 12,000 rpm for 20 minutes at 4°C. The supernatant was discarded, and the RNA pellet was washed with 500 μL of 70% ethanol. The solution was centrifuged at 7,500 rpm for 10 minutes at 4°C to pellet the RNA. The pellet was air-dried until the ethanol evaporated completely. Finally, the RNA pellet was resuspended in 20 μL of RNase-free water and incubated at 65°C before quantification using a NanoDrop spectrophotometer.

### cDNA preparation using Verso cDNA synthesis kit

cDNA was synthesized using the Verso cDNA Synthesis Kit (Thermo Fisher) according to the manufacturer’s guidelines. Briefly, 1 µg of RNA was placed in a PCR tube, and 1 µL of a primer mixture (mix) (comprising random hexamers and oligo-dT in a 3:1 ratio) was added to the RNA template. Nuclease-free water was added to bring the total volume to 12 µL. This mixture was incubated at 65°C for 5 minutes. After incubation, the mixture was cooled on ice, and an enzyme mixture (mix) containing the necessary buffer and components (8 µL) was added to the RNA-primer mix. The combined mixture (mix) was thoroughly mixed and then incubated in a dry bath at 45°C for 1 hour. The reaction was terminated by incubating the mixture at 95°C for 2 minutes. The synthesized cDNA was stored at -20°C for further use.

### RT-qPCR analysis

RT-qPCR was employed to perform real-time quantification of amplified genes using their specific primer sets. cDNA was synthesized from RNA served as the template, and a Sybr mix containing dNTPs, enzyme, buffer, and Sybr Green dye was added to the reaction mixture along with the template and primers. The reaction protocol included an initial denaturation step at 95°C for 10 minutes, followed by 35 cycles of denaturation at 95°C for 30 seconds, annealing at 60°C for 30 seconds, and extension at 72°C for 30 seconds. The results were analyzed and presented as relative fold changes compared to the expression level of a housekeeping gene (55).

### Immunofluorescence assay

Huh7 (mock) and HCV-infected cells, were grown on 18 mm glass coverslips at various time points post-infection. The cells were then fixed with 4% paraformaldehyde for 15 minutes at room temperature, followed by washing three times with 1x PBS. Permeabilization was attained by incubating the cells with 0.1% Triton X-100 in 1% PBS for 10 minutes at room temperature, followed by another set of three washes with 1x PBS. To block non-specific binding sites, the cells were incubated in 1% BSA in 1x PBS for 30 minutes at room temperature. For primary antibody incubation, the coverslips were treated with various primary antibodies, cGAS, STING, MAVS, TFAM, TOM20, PD1, RIG-I, HCV core and anti-dsRNA, and all prepared in 1% BSA in 1x PBS at the recommended dilutions. This incubation lasted for 1 hour at room temperature. Excess primary antibody was removed by washing the coverslips three times with 1x PBS for 5 minutes each. Next, the coverslips were incubated with an Alexa Flour-594 conjugated goat-anti-mouse secondary antibody, Alexa Flour-647 conjugated goat-anti-rabbit secondary antibody prepared in 1% BSA in 1x PBS, for 1 hour at room temperature in the dark. Following secondary antibody incubation, the coverslips were again washed three times with 1x PBS for 5 minutes each to remove any unbound secondary antibody. The nuclei were stained using DAPI solution and coverslips were then mounted on glass slides using a suitable mounting medium and sealed with nail polish to prevent drying and movement. Finally, the stained cells were spotted under a Zeiss Axio Observer 7 immunofluorescence microscope, and images were captured at appropriate magnifications to analyze the localization and expression of the target proteins. All antibody incubation steps were performed in the dark to prevent photobleaching, and gentle rocking or shaking was used during incubations to ensure even distribution of solutions (18).

### Western blotting

Total cellular protein lysates were prepared from mock- and HCV-infected Huh7 cells at various time points post-infection. Briefly, media was aspirated, and cells were washed with cold 1xPBS, thoroughly scraped, and centrifuged at 13,000× g for 5 minutes. After discarding the supernatant, the cell pellets were lysed using RIPA buffer (50 mM Tris, pH 7.5, 150 mM NaCl, 1% NP-40, 0.5% sodium deoxycholate, 0.1% SDS, and 1% protease inhibitor cocktail) and kept on ice for 30 minutes. The lysed cells were then centrifuged at 13,000× g for 15 minutes at 4°C. The supernatants were collected, and Protein concentration was measured by Bradford method using the Bradford reagent. Equal amounts of protein were loaded onto a 12% SDS-PAGE gel, followed by transfer to a PVDF membrane. The membrane was blocked with 5% skimmed milk in TBST for 1 hour, and then washed with TBST. The membrane was incubated overnight with specific primary antibodies, followed by 1-hour incubation with horseradish peroxidase-conjugated secondary antibodies. The specific protein bands were detected using enhanced chemiluminescence (ECL) substrate solution, which reacts with HRP. The resulting signal was captured on x-ray film for further analysis (56).

### Immunoprecipatation Assay

For the immunoprecipitation (IP) assay, harvested Huh7 cells were lysed using an IP lysis buffer consisting of 25 mM Tris-HCl (pH 7.5), 150 mM NaCl, 1% NP40, 2 mM EDTA, 10% glycerol, and a protease inhibitor mixture. Lysates were pre-cleared by incubating with Protein agarose beads to reduce non-specific binding. After pre-clearing, 150 to 200 μg of whole cell lysates were left to incubate overnight with the primary antibodies at 4°C. The antibody-protein complexes were then captured using Protein G-sepharose beads (G Biosciences India) and washed three times with IP lysis buffer to remove any non-specifically bound proteins. The immune complexes were eluted by adding SDS sample buffer and boiling the mixture. The eluted proteins were separated by SDS-PAGE, transferred to a PVDF membrane, and blocked with 5% skimmed milk in TBST. The membrane was probed with specific primary antibodies followed by HRP-conjugated secondary antibodies. Detection was performed using an enhanced chemiluminescence (ECL) system. The blots were developed on X-ray film to visualize the protein bands (57).

### Transfection of siRNA

Huh7 cells were transfected with siRNA targeting cGAS (#129125) STING (#128591) and RIG-I genes (#134222), as well as a non-coding siRNA as a control, using RNAiMax lipofectamine (#13778075) purchased from Thermofisher. Cells were seeded at a density of 0.8 × 10^6 cells per 60 mm culture dish and allowed to grow overnight. Prior to transfection, cells were incubated in OptiMEM media for 1 hour. For the transfection mixture, 200 pmol of siRNA was combined with 500 µL of OptiMEM media in a microcentrifuge tube and incubated at room temperature for 10 minutes. In another tube, 12 µL of lipofectamine was mixed with 500 µL of OptiMEM and also incubated at room temperature for 10 minutes. The two solutions were then combined and incubated for an additional 20 minutes at room temperature. The existing media was removed from the cells, and the siRNA-lipofectamine mixture was added dropwise to the cells in the 60 mm dishes. The cells were then incubated in a CO_2_ incubator for 6 hours. After this incubation, the transfection mixture was removed and replaced with complete media. Forty-eight hours post-transfection, cells were harvested for total RNA using TRIzol reagent. Then RNA was stored at -80°C for further use.

### cGAMP detection

The 2’3’-cGAMP ELISA was performed using the Cayman Chemical 2’3’-cGAMP ELISA Kit (Cat no. 501700) according to the manufacturer’s protocol. Total cellular protein lysates were prepared from mock- and HCV-infected Huh7 cells at various time points of infection. 2’3’-cGAMP detection using the ELISA kit involves several key steps. Initially, buffers are prepared by diluting immunoassay buffer C concentrate (10X) and wash buffer concentrate (400X) with ultrapure water. Samples, which can include cell lysates and supernatant are prepared by ensuring they fall within the standard curve’s linear range. The assay starts by preparing standards through serial dilution. The plate setup involves adding standards, samples, and controls, followed by the addition of 2’3’-cGAMP-HRP Tracer and Polyclonal Antiserum to each well. After 1 hour of incubation at room temperature with shaking, the plate is washed five times. TMB substrate solution is then added, followed by 30-minute incubation in the dark. The reaction is stopped with HRP stop solution, and the absorbance is measured at 450 nm. Data analysis includes constructing a standard curve to determine the 2’3’-cGAMP concentrations in the samples. Proper temperature control and gentle reagent handling are emphasized throughout the process (58).

## Funding

J.I. acknowledges the support of Ramalingaswami Fellowship Grant (BT/RLF/Re-entry/09/2015) from Department of Biotechnology (DBT), Ministry of Science and Technology, Govt. of India.

## Conflicts of Interest

Authors have declared no conflict of interest.

## Notes

### Competing Interest Statement

The authors have declared no competing interest.

