## Supplementary Figures for "Activation of cGAS-STING signaling pathway during HCV infection"

**Supplementary Data**

a

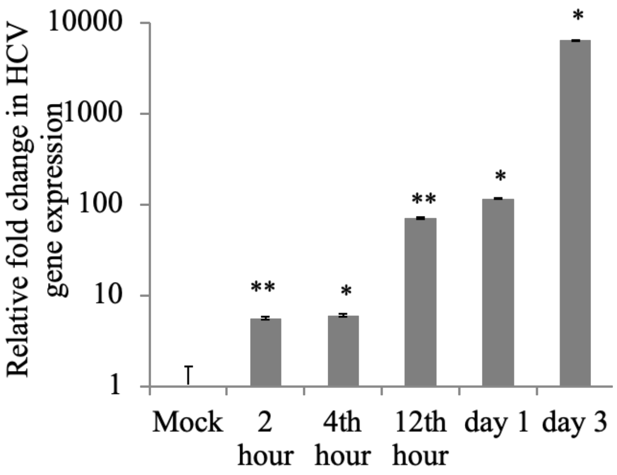


b

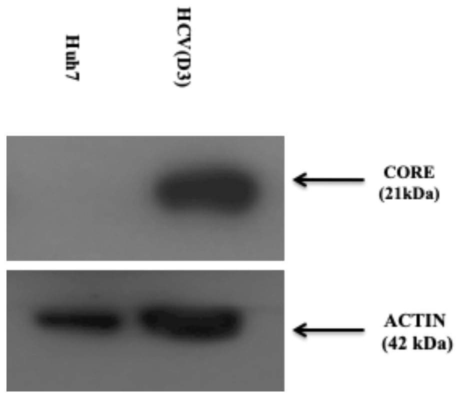


**Supplementary Fig. 1. Confirmation of HCV infection in human hepatoma cells**: (a) RT-qPCR image showing fold change in HCV gene expression at different time point of HCV infection compared to mock cells. (b) Western blot image showing HCV core protein expression (21kDa) in infected cells against mock infected cells. Actin (42kDa) shows the internal loading control. The values represent the means ± SDs from two independent experiments performed in duplicate. * p-value ≤ 0.05, ** p-value ≤0.01, for mock vs infection.

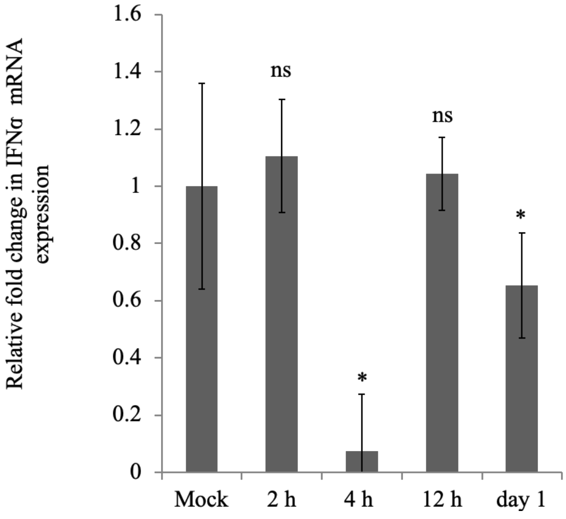


**Supplementary Fig. 2 Relative fold change in IFNα gene expression during HCV infection.** RT-qPCR image showing fold change in IFNα gene expression at different time point of HCV infection compared to mock cells. The values represent the means ± SDs from two independent experiments performed in duplicate. * p-value ≤ 0.05, ** p-value ≤0.01, for mock vs infection.
